# Choroid plexus defects in Down syndrome brain organoids enhance neurotropism of SARS-CoV-2

**DOI:** 10.1101/2023.06.12.544552

**Authors:** Mohammed R. Shaker, Andrii Slonchak, Bahaa Al-mhanawi, Sean D. Morrison, Julian D. J. Sng, Justin Cooper-White, Alexander A. Khromykh, Ernst J. Wolvetang

## Abstract

Why individuals with Down Syndrome (DS, trisomy 21) are particularly susceptible to SARS CoV-2 induced neuropathology remains largely unclear. Since the choroid plexus (CP) performs important barrier and immune-interface functions, secretes the cerebrospinal fluid and strongly expresses the ACE2 receptor and the chromosome 21 encoded TMPRSS2 protease, we hypothesized that the CP could play a role in establishing SARS-CoV-2 infection in the brain. To investigate the role of the choroid plexus in SARS-CoV-2 central nervous system infection in DS, we established a new type of brain organoid from DS and isogenic euploid control iPSC that consists of a core of appropriately patterned functional cortical neuronal cell types that is surrounded by a patent and functional choroid plexus (CPCOs). Remarkably, DS-CPCOs not only recapitulated abnormal features of DS cortical development but also revealed defects in ciliogenesis and epithelial cell polarity of the developing choroid plexus. We next demonstrate that the choroid plexus layer facilitates SARS-CoV-2 replication and infection of cortical neuronal cells, and that this is increased in DS-CPCOs. We further show that inhibition of TMPRSS2 and Furin activity inhibits SARS-CoV-2 replication in DS CPCOs to the level observed in euploid organoids. We conclude that CPCOs are a useful model for dissecting the role of the choroid plexus in euploid and DS forebrain development and enables screening for therapeutics that can inhibit SARS-CoV-2 induced neuro-pathogenesis.

## INTRODUCTION

The choroid plexus (CP) is a highly vascularized secretory tissue located within each ventricle of the vertebrate brain ^1^. The CP supports the central nervous system by producing and transporting cerebrospinal fluid (CSF) as well as a variety of signaling factors that orchestrate cortical development and neurogenesis, while preventing the infiltration of immune cells into the central nervous system ^1^. During early development, the CP anlage and the directly adjacent cortical hem, an important brain organizer region, are co-specified and both secrete and respond to morphogens such as Notch, WNT and BMP ^2^. This developmental path ensures that the CP is always in close vicinity to the cerebral cortex and this anatomical juxtaposition between CP and the cortex persists throughout life ^3^. The human CP acquires barrier, secretory and transport capacities after two weeks of development via acquisition of tight junctions, and influences the patterning and migration of cortical/hippocampal progenitors and Cajal-Retzius cells in the developing cortex through morphogen gradients established by apical motile cilia projected into the CSF by CP epithelial cells ^1^. To date, mammalian CP development has been predominantly studied in animal models ^4–6^ and it remains largely unclear to what extent these developmental processes are conserved in human. To model the human CP, several 3D culture systems have been developed that allow the generation of CP-like structures *in vitro* starting from human ESC-derived neuroepithelial cells ^7–10^.

Down syndrome (DS) is a genomic disorder with an incidence of 1 in 700 to 1000 live births ^11^ that is caused by the presence of a supernumerary chromosome 21 (HSA21). The extra copy of HSA21 (trisomy 21) results in neuropathological changes such as disorganized cortical lamination ^12^, altered cerebellar organization and function ^13^, and a hypocellular hippocampal dentate gyrus ^14^. The DS cerebral cortex further exhibits a reduction in excitatory neurons ^15^, an increased production of astrocytes and inhibitory neurons ^16^, as well as defective oligodendrocyte differentiation and myelination ^17^. The developing DS brain and a DS mouse model ^18^ further display ventriculomegaly that was linked to increased dosage of the HSA21 genes *PCNT* and *PCP4* involved in cilia function, and defective cilia in human DS fibroblast cells were previously reported ^19^. Previously cerebral organoids derived from DS iPSC were found to recapitulate various aspects of altered DS brain development ^20^, including defective generation of cortical neurons ^21^, excessive interneurons production ^20^, and Alzheimer’s disease (AD)-like pathology ^22^. Accumulating clinical evidence indicates that SARS-CoV-2 infection of people with DS is associated with a 4-fold increased risk in hospitalization and a 10-fold increased risk of death as compared to euploid counterparts that cannot be readily explained by comorbidities ^23, 24^. It was hypothesized that increased susceptibility to COVID-19 pathology may in part be explained by an exaggerated interferon response brought about the increased gene dosage of interferon pathway genes located on HSA21 and in part by defects in systemic immune system function ^25, 26^ previously linked to increased bacterial and viral infections in people with DS ^27–29^. It is also possible that the increased dosage of TMPRSS2, a HSA21 gene that codes for a protease that promotes interaction between SARS-CoV-2 spike protein and the ACE2 receptor ^30^, plays a role. Neurotropism of SARS-CoV-2 is increasingly recognized as a possible driver of long-term cognitive and sensory impairment (long-COVID) ^31^. However, since people with DS intrinsically exhibit a range of progressive inter-individually highly variable cognitive deficits and a dramatically increased risk of early onset Alzheimer’s-like disease, the long-term impact of SARS-CoV-2 infection on cognitive function of DS people is difficult to quantify. It also remains to be determined to what extent vertical transmission of SARS-CoV-2 from mother to fetus can interfere with brain development, and whether this more severely impacts DS brain development.

Here we report the generation of human cortical brain organoids that are both surrounded by a functional CP and contain properly patterned and developing cortical cell types (CPCOs). We demonstrate that these CPCOs display typical neuropathological changes of DS such as an imbalanced production of excitatory and inhibitory neurons and reduction in oligodendrocyte progenitor cells (OPCs). We further discovered that DS CPCOs exhibit aberrant ciliogenesis and defective polarity of the CP. Strikingly, we show that the CP compartment of CPCOs strongly facilitates the neuroinvasion and neurotropism of SARS CoV-2 and that this is increased in DS. We further show that treatments with FDA approved drugs that inhibit TMPRSS2 activity as well as a Furin inhibitor and Remdesivir reduce SARS CoV-2 replication in DS organoids to a level comparable to the euploid group, suggesting that increased gene dosage of TMPRSS2 in the DS CP may be involved, and indicating that CPCOs are a suitable model for identifying drugs that can reduce the impact of SARS-CoV-2 on the mature and developing CNS.

## RESULTS

### Self-Assembly of Choroid Plexus Organoids Recapitulates Embryonic Development

Human neuroectodermal (hNEct) are primed to exclusively develop into tissues of the anterior body ^32^ and are fated to form the cortex dorsally and the cortical hem ventrally, which next differentiates into the CP. We therefore reasoned that hNEct cells would be an appropriate starting population for the generation of self-organizing cortical organoids surrounded by ventricular structures derived from the CP (here termed CP-Cortical organoids (CPCOs)). We exposed human ES and iPSC lines (Figure S1A) (H9 ^33^, WTC ^34^ and G22 ^34^) to dual SMAD inhibition with SB and LDN for 3 days, which resulted in the efficient generation of hNEct cells that generate neural stem cells characterized by the expression of SOX2 and NESTIN (Figure S1B). hNEct cells were next lifted to form 3D aggregates on ultra-low attachment 6 well plates in N2 medium supplemented with bFGF, resulting in the formation of spheres with a neuroepithelial layer (Figure 1A, day 7) and multiple SOX2 expressing rosettes at the center (Figure S1C). To mimic the secretion of BMP4 and WNTs by the cortical hem that specifies the dorsal cortical hem into CP *in vivo* ^8^, we treated these spheres with BMP4 and CHIR99021 to promote CP formation (Figures 1A, S1C). Since the amount of BMP4 signaling is known to promote CP lineage differentiation at the expense of neural lineages ^9, 10^, we exposed hNEct to increasing BMP4 dosages during the initial 4 days of the CPCO protocol. Low doses of BMP4 (2.5 ng) reduced the proportion of SOX2 expressing cells in the early organoids to 50% in an outside-inside fashion and was further adopted as the optimal concentration (Figure S1D), and increasing the dosage to 50 ng/ml BMP4 further reduced the proportion of SOX2 expressing cells, as expected (Figure S1D). Quantification of mRNA levels of cortical hem (*MSX1/2*) and CP genes (*AQP1*, *TTR*, and *KLOTHO*) by quantitative PCR (qPCR) revealed that 21 days of CHIR99021 and BMP4 treatment resulted in a significant induction of *MSX1/2* expression followed by a sharp reduction from day 28 to day 84 (Figure 1C), suggesting the induction of cortical hem. In agreement with these data, cross-sectioned organoids revealed co-expression of cortical hem markers MSX1/2 and LMX1A proteins in the folded epithelial cells by day 14 (Figure 1D). Between day 21 to day 28, we detected the emergence of thin epithelial layers surrounding the organoid (Figure 1A), and qPCR demonstrated the concomitant upregulation of CP markers, *AQP1*, *TTR* and *KLOTHO* (Figure 1C). Consistent with these observations, immunostaining revealed the expression of the definitive CP markers TTR and LMX1A proteins in these epithelial layers (Figure 1D). Organoid size gradually increased over the first 28 days (Figure 1B), until reaching a mean core diameter size of 1.2 mm (Figure 1B), and a final diameter of 1.9 mm at day 56 and day 120 (Figure 1B). High-resolution 3D imaging identified multiple CPs emerging from a single organoid that formed an intact epithelial covering of the entire organoid as indicated by the tight junction marker ZO1 (Figure 1E). These cells also robustly expressed KLOTHO, an anti-aging protein known to be expressed in mouse CP ^33^, confirming that KLOTHO expression in the CP is evolutionary conserved and suggesting that its role in these cells can be studied with these human CP organoids. To exemplify the reproducibility of the system, we generated CPCOs from different hiPSC lines (Figures S2A, S2B), and found that more than 77% of the organoids exhibited the characteristic thin TTR^+^ epithelial layers enveloping the organoid, independent of cell line, clone, or batch (Figure S2C). We further found that these organoids can survive for prolonged periods (currently 8 months in culture) without obvious signs of deterioration. Collectively, our data demonstrate that this protocol recapitulates the *in vivo* developmental stages of cortical hem patterning and CP tissue formation, and outlines a rapid protocol for generating human brain organoids that are encased in CP.

**Figure 1.**
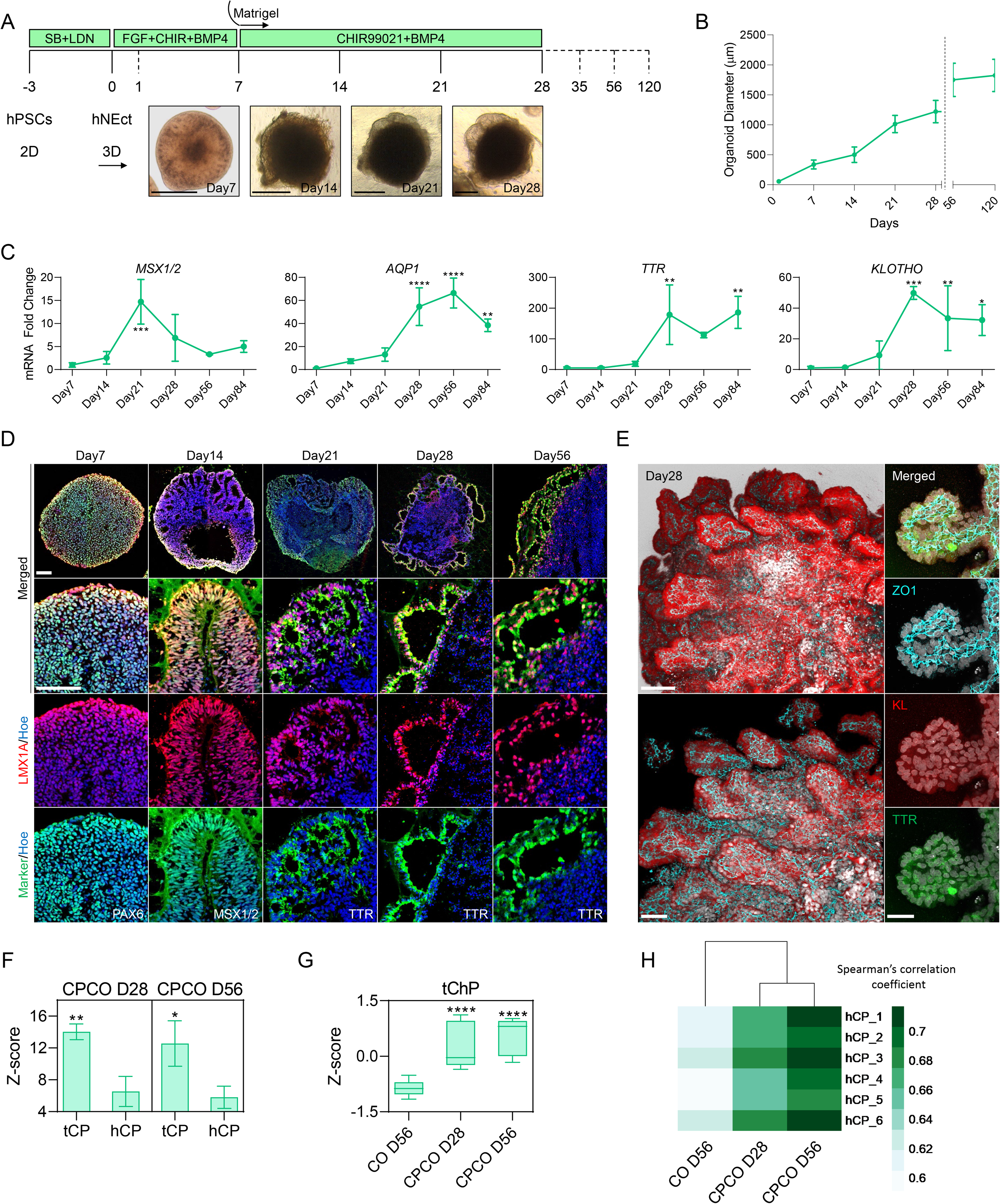
Generation of Human Self-organizing multiple CPCOs in 3D from Neuroectoderm. (**A**) Schematic representation of the strategy used to generate CPCOs from hPSCs. Below brightfield images showing the developmental stages of CPCO overtime *in vitro*. Scale bar = 500 µm. (**B**) Graph showing the growth (average diameter) of CPCOs at different stages of *in vitro* culture. Data are presented as mean ± standard deviation (n = 4). (**C**) qRT-PCR of cortical hem marker (*MSX1/2*) and CPs definitive markers (*AQP1*, *TTR* and *KLOTHO*). All values were normalized to GAPDH levels of their respective samples and expressed relative to Day 7 values to obtain the fold change. Data are shown as mean ± standard deviation; Number of independent experiments = 4. *p < 0.05, **p < 0.01, ***p < 0.001, ****p < 0.0001 via one-way ANOVA. (**D**) Analysis of immunostaining of sections organoids at days 7, 14, 21, 28 and 56 showing the proteins expression of neuroectoderm markers PAX6 (Green) and LMX1A (Red), cortical hem markers MSX1/2 (Green) and LMX1A (Red), and CP definitive markers TTR (Green), LMX1A (Red). All sections were counterstained with Hoechst 33342 (Blue). Scale bar = 72 µm. (**E**) Wholemount immunostaining images of CPCO at day 42 of differentiation. Left images showing the multiple CPs formation at the periphery of an organoid stained with KLOTHO (Red) and ZO-1 (Cyan), Scale bar = 110 µm. Right images are 100X magnification of a single CP stained with ZO-1 (Cyan), KLOTHO (Red) and TTR (Green), Scale bar = 30 µm. (**F**) Bar graph representing the average z-score per column (pool of three replicates), showing distribution of marker genes for tCP and hCP obtained from bulk RNA-seq of CPCOs at days 28 and 56. Data representing the average z-score per column (pool of three replicates). Data are presented as the mean ± standard deviation. *p < 0.05, **p < 0.01 via Student’s t-test. (**G**) Box blot showing distribution of marker genes for tCP obtained from bulk RNA-seq of CPCOs at days 28 and 56 and traditional “naked” COs at day 56. Data are presented as minimum to maximum, with notches are centered on the median. ****p < 0.0001 via one-way ANOVA. See Fig. S2E for individual genes used. (**H**) Heatmap comparing the Spearman correlation coefficient of the bulk RNA-seq of CPCOs at days 28 and 56, and traditional “naked” COs at day 56 to adult human CP ^99^.

In an effort to further simplify the protocol, we attempted to generate CPCOs without embedding hNEct spheres into the Matrigel. However, non-embedded organoids treated with BMP4 and CHIR99021 for 28 days failed to generate the thin epithelial layers surrounding the organoids and led to substantial amounts of cell death in the culture (Figure S1E), indicating that ECM plays a critical role in CP morphogenesis *in vitro*.

We next compared the bulk RNA transcriptomes of day 28 and day 56 CPCOs with “naked” (non-BMP4 and CHIR99021 treated) cortical organoids (COs). This revealed that the expression of cortical hem and CP markers in day 28 and day 56 CPCOs generally increased over time, suggesting the development of CP is progressing beyond 28 days of *in vitro* culture, and was significantly higher than those observed in “naked” COs at day 56 (Figure S2D). This data is in agreement with our observation that the CP in day 56 CPCOs consists of multiple epithelial layers (TTR^+^ and LMX1A^+^) whereas only a single thin discontinuous layer of CP is found in CPCOs at day 28 (Figure 1D). *In vivo*, BMP and SHH gradients pattern the telencephalic CP (tCP) and hindbrain CP (hCP) along the dorsal axis of the neural tube, resulting in CPs with distinct transcriptome profiles ^35^. At days 28 and 56, our CPCOs showed a significant enrichment for genes related to tCP rather than hCP (Figures 1F, S2E), in agreement with the fact that our protocol involves treatment with BMP4 without addition of SHH, and these genes were again expressed at higher levels in CPCOs than in day 56 “naked” COs (Figure 1G). At a whole transcriptome level, CPCOs, but not “naked” COs, showed a high correlation to adult human CP tissue (Figures 1H, S2F).

### Progressive Maturation of Choroid Plexus in CPCOs

To determine whether these self-organized CP layers in CPCOs exhibit similar properties to the *in vivo* CP, we next examined the establishment of epithelial polarity, abundance of mitochondria, ciliogenesis, and CSF secretion ^36^. We first assessed the establishment of epithelial polarity in the CP epithelium which is governed by tight junctions, gap junctions, adherent junctions, and basal lamina (Figure 2A). Bulk RNA-seq data identified a progressive enrichment of genes associated with these structures in CPCOs over time as compared to “naked” COs (Figures 2B, 2C, S3A). qPCR analysis revealed a gradual increase of selected tight junction genes such as *CLDN11* and *CLDN12* (Figure S3B), although *CLDN2* was only significantly increased by day 28 (Figure 2D). Similarly, mRNA levels of the gap junction gene *GJB2* and adherent junction genes *CDH5* and *PCDH18* also gradually increased (Figures 2D, S3B). Consistent with the widespread and contiguous expression of the tight junction marker ZO1 in the CP layers (Figure S3C), transmission electron microscopy (TEM) confirmed the formation of tight junctions as well as the extensive accumulation of mitochondria that is typical of CP cells (Figure 2E). The establishment of tight junctions is also critical for polarization of CP into distinct basal components ^37^, and we therefore examined whether the CP cells of CPCOs would exhibit apicobasal polarity of membrane proteins critical for normal CP epithelial cell function. Labeling CPCOs with ZO1 and LAMININ, which accumulate on the apical and basal side of polarized CP epithelium, respectively, revealed the correct formation of the basal membrane and apical polarity (Figure 2F). We reasoned that the establishment of CP epithelial polarity in CPCOs, as well as the increase of mitochondria in CP that is required for CP homeostasis, cerebrospinal fluid secretion, neurotrophic factor transport, and barrier function, was an indication that the CPs in our CPCOs should have the potential for CSF production and effective barrier formation. RNA-seq analysis indeed confirmed the significant enrichment of genes that code for CP secreted proteins in CPCOs at day 28 compared to “naked” COs at day 56 (Figure 2G) and revealed that their expression significantly increases as CPCOs further develop (Figure 2G, CPCOs day 56). We detected expression of *IGF2*, a growth factor known to be secreted in the CSF and that stimulates ependymal NSCs proliferation ^38^, ECMs genes such as *SPARC* that are involved in uptake and delivery of proteins from blood to the CSF ^39^, as well as a number of enzymes, secreted proteins, and membrane proteins, that are all involved in CSF generation and secretion (Figure 2H). Furthermore, staining showed the correct apical localization of the water channel aquaporin 1 (AQP1) in the CP of CPCOs at day 28, a protein involved in CSF secretion (Figure 2I). We next used a multiplex ELISA to determine the CPCO secretome released in the culture medium. This revealed a significant increase of CST3 and B2M of CPCOs as compared to “naked” COs (Figure 2J). The abundance of APP secretion was however similar between CPCOs and “Naked” COs (Figure 2J).

**Figure 2.**
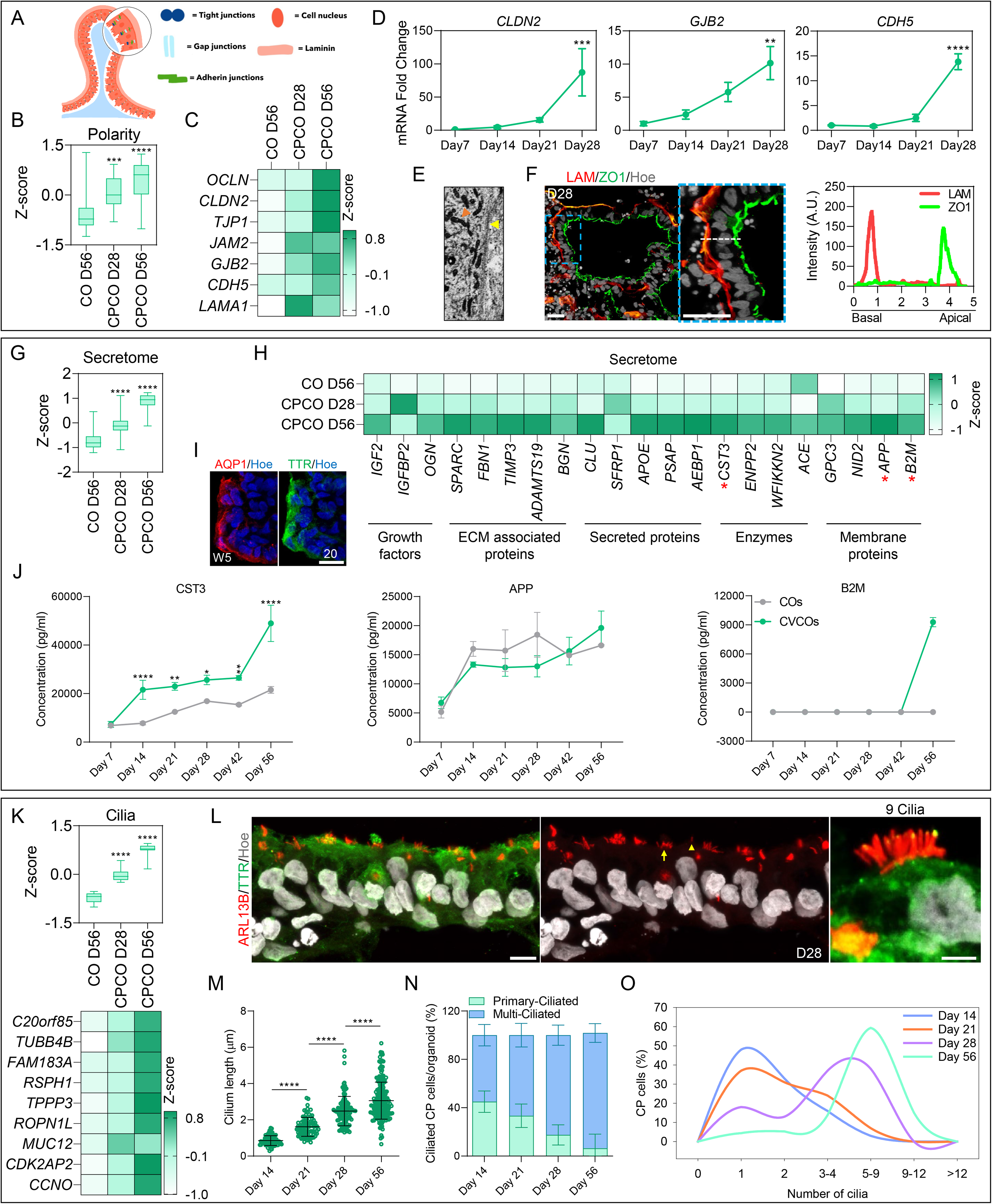
Development and Functional Analysis of CP in CPCOs. (**A**) Schematic diagram outlining the apical-basal polarity of the CP. (**B**) Box blot showing distribution of polarity marker genes (listed in **C,** and Fig. **S3A**) obtained from bulk RNA-seq of CPCOs at days 28 and 56 and traditional “naked” COs at day 56. Data are presented as minimum to maximum, with notches centered on the median. ***p < 0.001, ****p < 0.0001 via one-way ANOVA. (**C**) Heatmap expression of representative marker genes related to apicobasal polarity within the bulk RNA transcriptomes of CPCOs at days 28 and 56 and traditional “naked” COs at day 56. Values are shown as z-score. (**D**) qRT-PCR of marker genes related to apicobasal polarity (*CLDN2, GJB2, and CDH5*) in CPCOs. All values were normalized to GAPDH levels of their respective samples and expressed relative to Day 7 values to obtain the fold change. Data are shown as the mean ± standard deviation; **p < 0.01, ***p < 0.001, ****p < 0.0001 via one-way ANOVA. Number of independent experiments = 3. (**E**) Transmission electron microscopy of CPCOs at day 28 showing the high-density of mitochondria (orange arrowhead) and tight junction formation (yellow arrow). (**F**) Analysis of immunostaining of CPCO sections at day 28 with the basal polarity marker LAMININ (Red) and apical marker ZO1 (Green). The section was counterstained with Hoechst 33342 (Grey). Scale bar = 20 µm. Dotted white line represents the average intensity of LAMININ and ZO1 expression along the apicobasal of CP epithelium plotted in the graph. (**G**) Box blot showing distribution of marker genes (listed in **H**) associated with CSF secretome obtained from bulk RNA-seq of CPCOs at days 28 and 56 and traditional “naked” COs at day 56. Data are presented as minimum to maximum, with notches are centered on the median. ****p < 0.0001 via one-way ANOVA. (**H**) Heatmap expression of marker genes related to CSF secretome within the bulk RNA transcriptomes of CPCOs at days 28 and 56 and traditional “naked” COs at day 56. Values are shown as z-score. Red asterisks represent the genes selected for ELISA experiment shown in panel (J). (**I**) Analysis of immunostained CPCO sections at day 35 showing protein expression of water channel aquaporin 1 (AQP1) in Red, and TTR in Green in CP. The section was counterstained with Hoechst 33342 (Blue). Scale bar = 20 µm. (**J**) Luminex multiplex/ELISA showing CSF secretome protein markers in media of CPCOs at days 7 14, 21, 28, 42, and 56. Data are shown as mean ± standard deviation; Number of independent experiments = 3. *p < 0.05, **p < 0.01, ****p < 0.0001 via one-way ANOVA. (**K**) Box blot showing distribution of marker genes (listed below in the heatmap) associated with cilia obtained from bulk RNA-seq of CPCOs at days 28 and 56 and traditional “naked” COs at day 56. Data are presented as minimum to maximum, with notches are centered on the median. ****p < 0.0001 via one-way ANOVA. Below heatmap expression of marker genes related to cilia. Values are shown as z-score. (**L**) Analysis of immunostained CPCO sections at day 28 showing ARL13B (Red) protein expression in cilia of CP marked with TTR (Green). The section was counterstained with Hoechst 33342 (Grey). Scale bar = 20 µm., magnified image scale bar = 5 µm. Yellow arrow indicates a multi-ciliated cell, yellow arrowhead indicates a mono-ciliated cell. (**M**) Quantification of cilia length in CPCOs at days 1, 21, 28, and 56 of differentiation. Data are presented as mean ± standard deviation; **** P<0.0001 via One Way ANOVA Number of independent experiments = 3. Individual dots represent a cilium length. (**N**) Stacked bar graph showing the percentage of cells with a single cilium and multiple cilia in human CP cells in CPCOs at days 7, 21, 28, and 56 of differentiation. Data are presented as mean ± standard deviation. Number of independent experiments = 3. (**O**) The distribution of cilia length in CPCOs culture for days 14, 21, 28, and 56. The data is presented as a percentage of CP cells with a single primary cilium or multi-cilia.

Primary cilia in the CP are essential for regulating the flow and transport of CSF and can be characterized according to their length, motility and number per cell ^40^. *In vivo*, the CP consists mainly of epithelial cells that are multi-ciliated, with tufts of cilia ranging from 4-8 cilia per cells, but also contains a small fraction of CP cells implicated in chemo- and/or osmosensation that extend one primary cilium into the CSF ^41^. How and when these cell types are specified in human embryos remains largely unclear. Since the generation of CPCOs closely mimics the progressive temporal morphogenic sequence of events of normal CP development *in vivo* (Figure 1), we examined ciliogenesis in our human CPCOs cultured for 14, 21, 28, and 56 days. Bulk RNA-seq confirmed the significant and progressive enrichment of cilia associated genes in CPCOs as compared to COs (Figure 2K). Immunostaining for cilia with the ARL13B antibody demonstrated that human CP cells in CPCOs are ciliated (Figure 2L), and that cilia length significantly increases over time, with a mean value of 0.7µm, 1.6µm, 2.3µm, and 3µm at day 14, day 21, day 28, and day 56, respectively (Figure 2M). At day 14, the developing CP contains almost equal amounts of mono- and multi-ciliated TTR expressing cells, and then displays a progressive increase in multi-ciliated cells to approximately 66% at day 21, 82% at day 28, and 95.4% at day 56 that is accompanied by a concomitant decrease in mono-ciliated CP cells (Figure 2N). The number of cilia on CP cells in mammals has been reported to range from 4-8 cilia per cell in rats to 50 cilia per cell in salamander ^1^. Our data show that human CP cells in CPCOs display a distinct shift in the number of cilia per cell over time in culture between day 14 and day 56, at which point >50% of all CP cells possess between 5 and 9 cilia per cell (Figure 2O). Collectively, these findings indicate that the self-assembled CPCOs recapitulate key aspects of *in vivo* CP ciliogenesis, suggesting that CPCOs may provide a useful model for investigating diseases such as Down syndrome ^18^ and Bardet-Biedl syndrome ^42^ in which defective primary cilia are thought to cause brain ventriculomegaly and hydrocephalus, respectively.

### Specialized Functional Cellular Compartments of the Cerebral Cortex Arise in CPCOs

In addition to the CPs components outlined above, CPCO organoids also possess other neural compartments. During embryonic development, the NEct-derived rostral neural tube gives rise to the CP in the dorsomedial telencephalon and cortical plate dorsally ^32^. We investigated whether these key cellular compartments of the cortical plate layers arise in our organoids in addition to the CP (Figure 3A) by examining the expression of different cortical neuronal markers known to be expressed in cortical organoids. We found a gradual reduction of *PAX6* mRNA over time (Figure 3B), and a concomitant increase in layer VI, V, IV, and II/III neuronal markers as indicated by the expression of *TBR1*, *CTIP2*, *SATB2*, and *CUX1/2*, respectively (Figure 3B). Staining for these neuronal proteins in midline sectioned organoids revealed specification of cortical TBR1^+^ and CTIP2^+^ neurons in the core of the organoid at day 28 (Figures 3C, S4A), whereas SATB2^+^ and CUX1^+^ neurons were only detected at day 56 of organoid maturation (Figure 3C). The specification of cortical layers occurred in close vicinity to the ventricle-like structures formed by the CP epithelial cells (Figures 3C, S4A). Astrocytes marked by GFAP were also specified in CPCOs and COs at day 56 of differentiation (Figures 3D), but not in CPCOs at day 28 (Figure S4C) in agreement with the notion that astrogliogenesis commences around 50 days ^43^. Furthermore, we detected oligodendrocytes marked by PDGFRA and CNPase at day 56 (Figure S3D), and co-localisation of MBP and neurofilament was observed in the mature organoids at day 150 (Figures 3E, S3D), suggesting the onset of myelination. This architectural structure of CP, ventricle-like structures and cortical neural cell types (Figure S4D) thus closely mimics the developing *in vivo* forebrain structure.

**Figure 3.**
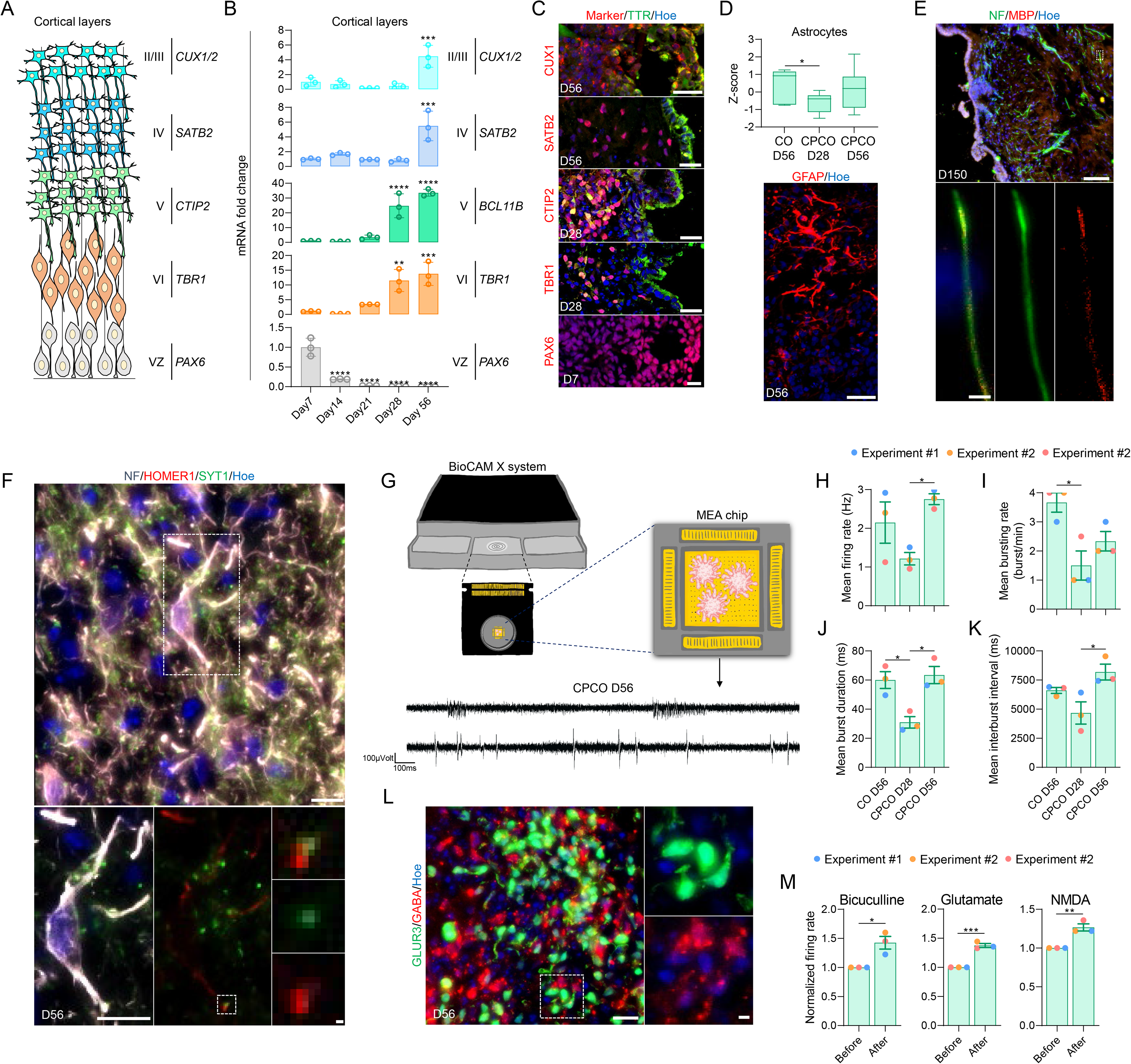
CPCOs contain functional cortical neurons with progress maturation over time in culture. (**A**) Schematic diagram of the development of the first four cortical plate layers: VZ is ventricle zone. (**B**) qRT-PCR of cortical ventricular zone (VZ) and cortical neuronal layer gene markers (*TBR1*, *CTIP2*, *SATB2* and *CUX1*/2). All values were normalized to GAPDH levels of their respective samples and expressed relative to Day 7 values to obtain the fold change. Data are shown as the mean ± standard deviation; **p < 0.01, ***p < 0.001, ****p < 0.0001 via one-way ANOVA. Number of independent experiments = 3. (**C**) Analysis of immunostaining of CPCO sections at days 7, 28, and 56, showing neural stem cells marked by PAX6 (Red, and cortical neurons in layers VI, V, IV, and II/III marked by TBR1 (Red), CTIP2 (Red), SATB2 (Red), and CUX1 (Red), respectively. All sections were counterstained with Hoechst 33342 (Blue). Scale bar = 10 µm for PAX6, Scale bar = 20 µm for TBR1, CTIP2, and SATB2, Scale bar = 30 µm for CUX1. (**D**) Box blot showing distribution of marker genes (listed in Fig. **S4B**) associated with astrocytes obtained from bulk RNA-seq of CPCOs at days 28 and 56 and traditional “naked” COs at day 56. Data are presented as minimum to maximum, with notches centered on the median. *p < 0.05 via one-way ANOVA. Below image shows CPCOs at day 56 immunostained with GFAP (Red). The section was counterstained with Hoechst 33342 (Blue). Scale bar = 30 µm. (**E**) Analysis of immunostaining of CPCO sections at days 150 showing myelinated neurons marked by Neurofilament (Green) and MBP (Red). Scale bar = 100 µm. Dotted white box indicates zoomed images below. Scale bar = 2 µm. (**F**) Analysis of immunostaining of CPCO sections at days 56 showing neurons with post- and pre-synapses marked by Neurofilament (Grey), HOMER1 (Red), and SYNT1 (Green). Scale bar = 10 µm. Dotted rectangular white box indicates zoomed images of a neuron showed below with scale bar = 1 µm. Doted square white box indicates zoomed imaged of connected pre- and post-synapse with scale bar = 1 µm. (**G**) Schematics of extracellular recordings from CPCOs on MEA. Below are the representative transient plots from neural activities recorded in CPCOs at day 56. Scale bar, 100 ms (horizontal), 100 µVolt (vertical). (**H-K**) Bar graphs show the changes in the patterns of neural activity in CPCOs at days 28 and 56 compared to traditional “naked” COs at day 56. Bar graphs represent mean ± SD in firing rate (**H**), mean bursting rate (**I**), mean burst duration (**J**), mean interburst interval (**K**). *p < 0.05 via one-way ANOVA. (**L**) Magnified image of sectioned CPCOs at day 56 immunostained for glutamatergic (GLUR3, Green) and GABAergic (GABA, Red) neurons on day 56. All sections were counterstained with Hoechst 33342 (Blue). Scale bar = 20 µm, scale bar for magnified image = 10 µm. (**M**) Bar graphs showing the changes in the patterns of mean firing rate before and after drug treatments. The following drugs were used: 50 µM Glutamate, 10 µM NMDAA, and 50 µM Bicuculline. *p < 0.05, **p < 0.01, ***p < 0.001 via Student’s t-test.

To evaluate the functional properties of the cortical neurons, we first confirmed that mature neurons exhibited the expression and juxtaposition of pre- and post-synaptic markers SYT1 and HOMER1, respectively (Figure 3F), as morphological evidence of synaptic contacts. We then characterized key electrophysiological parameters using high density multielectrode arrays (MEAs) (Figure 3G). MEA analysis detected spontaneous electrical activity in CPCOs from day 28 (an example of the traces at day 56 is shown in Figure 3G). Over time, the spontaneous neural firing/bursting rates progressively increased (Figures 3H-K). Conspicuously, firing rates increased significantly at day 56, when GFAP^+^ cells were observed in the CPCOs (Figure 3D). However, the burst rate did not significantly increase in CPCOs at day 56 in contrast to “naked” COs (Figure 3I). We speculate that this difference may be due to the different proportions of neuronal subtypes present in these two types of organoids. We indeed noted that 56 day-old CPCOs contained glutamatergic (GLUR3^+^) and GABAergic (GABA^+^) neurons that were located adjacent to each other (Figure 3L), consistent with the specification of GABAergic inhibitory neurons during *in vivo* cortical development ^44^. Treatment with the GABA receptor antagonist bicuculline (50µM) significantly increased the firing rate in CPCOs (Figures 3M, S4E), confirming the presence of inhibitory synaptic transmission, while acute glutamate (200µM) and NMDA (50µM) treatments significantly increased the firing rate (Figures 3M, S4E), demonstrating the presence of active glutamatergic and GABAergic neurons. Collectively, these data suggest that neural networks with excitatory and inhibitory neural circuits were well established within the CPCOs at day 56.

### Down syndrome CPCOs recapitulate key aspects of DS brain pathology

DS is caused by trisomy 21, and leads to developmental delay and intellectual defects ^45^. Several DS mouse models previously showed evidence of altered production of excitatory and inhibitory forebrain neurons in the neocortex ^45, 46^. Both the human developing DS brain and Ts65Dn DS mice further display defects in oligodendrocyte differentiation and myelination ^17^. Moreover, trisomy 21 fibroblasts exhibit a reduction in cilia formation and function ^19^, making DS a particularly interesting condition to model in CPCOs. We therefore generated CPCO organoids from an iPSCs line derived from patient with DS (DS18) and its isogenic euploid counterpart, the iPSCs line (EU79) ^47^ (Figure 4A). The size of the DS CPCO organoids was comparable to the euploid organoids (Figure S5A), suggesting similar growth rates. Quantification of the percentage of CP and cerebral cortex cell populations in at least 3 different batches of CPCO organoids generated from the EU79 and DS18 lines revealed that CPCOs of euploid and DS lines both reproducibly organized into CP and cerebral cortex regions with similar cell proportions across different batches (Figures 4B, 4C). To characterize the isogenic DS and euploid CPCOs in more detail, we performed RNA-seq analysis (three organoids per replicate, n = 3) on day 28 organoids (Figures 4D, S5B, Table S1). This revealed that euploid and DS organoids exhibited distinct transcriptional profiles (Figure S5B) with 962 genes with increased and 997 genes with decreased expression in DS compared to the isogenic euploid organoids (Figure 4D, Table S1). 22 genes (9.7%) located on HSA21 were among the top 500 differentially expressed genes (Figure S5D), five of these genes (MX2, TMPRSS2, ADAMTS5, and RUNX1) are associated with viral infection (Figure S5D). Enrichment analysis of upregulated genes in DS organoids revealed enrichment for many nervous system development, cilium movement and motility-related genes, as well as cell adhesion and basement membrane genes (Figure 4E), but no changes in CP or secretome associated genes (Figure S5C). DS organoids displayed a significant increase in the expression of a subset of ciliogenesis markers (Figure 4F). In contrast, immunostaining results of cilia with ARL13B antibody indicated that CP cells in DS CPCO organoids at 56 days are predominantly mono-ciliated (Figures 4G, 4H), and generated significantly less ciliated CP cells than the corresponding euploid CPCOs (Figure 4I), although the average cilium length was comparable between the two groups (Figure S6F). Unexpectedly, we also detected a significant alteration in apicobasal polarity marker genes (Figure S6A) and proteins (Figure S6C) in DS organoids. To further investigate possible defects in cell polarity, euploid and DS organoids were stained with ZO1 and β-catenin, two proteins involved in apical-basal epithelial cell polarity ^48^ (Figure 4J). In euploid organoids, these proteins correctly segregated to their respective compartments in CP, as expected (Figure 4J). However, in DS organoids derived from DS18 and G21 lines, ZO1 and β-catenin remained conspicuously co-localized (Figure 4J). We quantified the percentage of overlapping between ZO1 and β-catenin of CP epithelial cells ^48^. Quantification demonstrated that ZO1 and β- catenin overlapped in 14% and 9.4% of CP cells in CPCOs from euploid EU79 and G22 lines, but remained co-localised in 75% and 44.7% of CP cells in CPCOs from the Down syndrome DS18 and G21 lines (Figure 4L).

**Figure 4.**
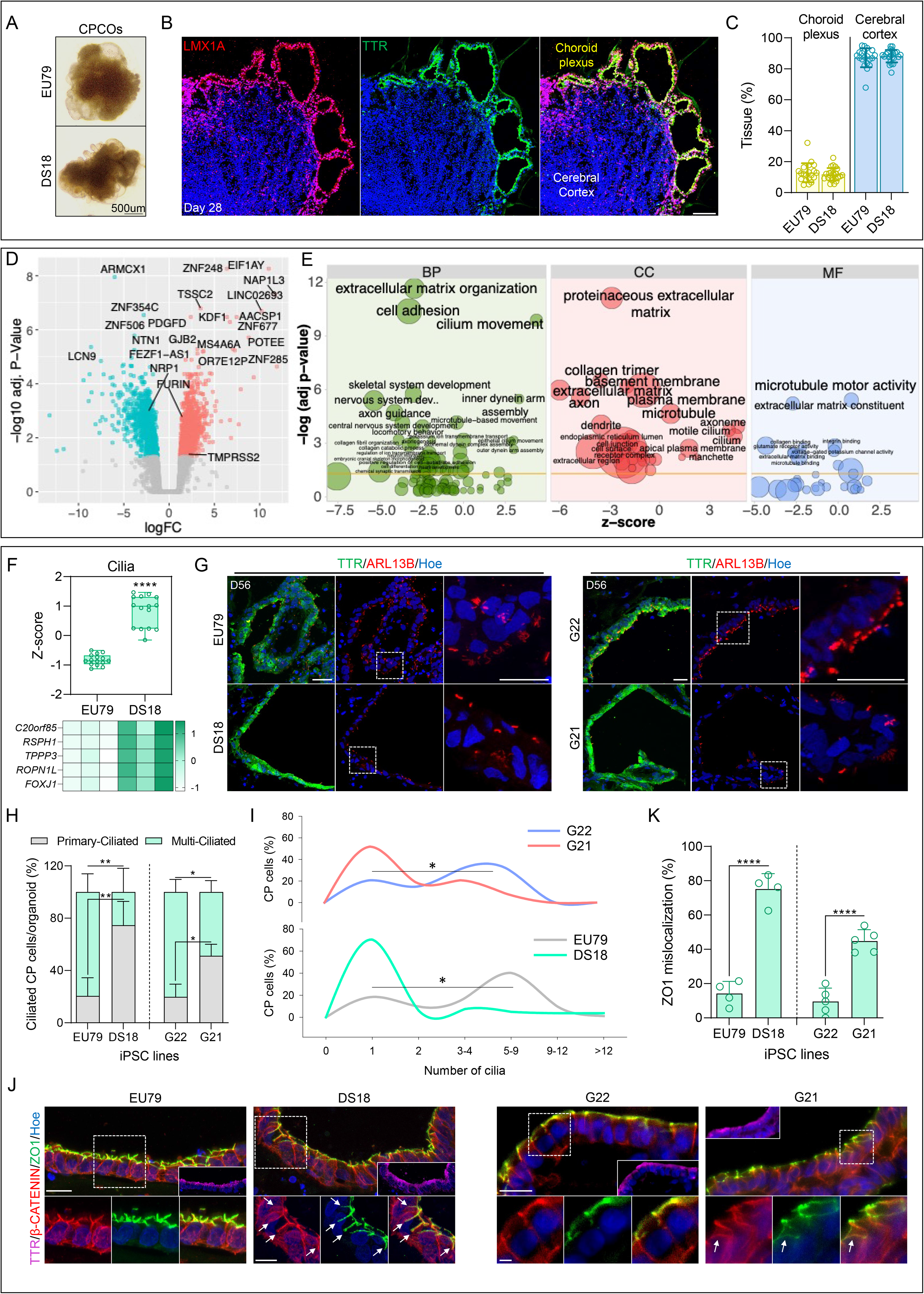
CPCOs modeling of Down syndrome. (**A**) Representative bright-field images of euploid and DS-iPSC derived CPCOs at day 28. Scale bar = 500 µm. (**B**) Representative image of euploid-iPSC derived CPCOs at day 28 immunostained with LMX1A (Red), and TTR (Green). The section was counterstained with Hoechst 33342 (Blue). Scale bar = 100 µm. (**C**) Quantification of choroid plexus and cerebral cortex tissues compartments in at least 3 different euploid (EU79) and DS (DS18) hiPSC derived CPCOs. Individual dots represent a single organoid. (**D**) Volcano plot highlighting differentially expressed genes in euploid and DS CPCOs at day 28. Significant up-regulated genes are shown in red and down-regulated genes are shown in cyan. Top-most differentially expressed genes as well as *TMPRSS2*, *FURIN*, *NRP1* genes are labelled. (**E**) Gene ontology (GO) enrichment analysis of differentially expressed genes (DEGs) in DS CPCOs compared to euploid organoids. Z-scores indicate the cumulative increase or decrease in expression of the genes associated with each term. Size of the bubbles is proportional to the number of DEGs associated with respective GO term. BP – biological processes, CC – cellular components, MF – molecular functions. (**F**) Box blot showing distribution of marker genes (listed below in the heatmap) associated with cilia obtained from bulk RNA-seq of CPCOs at day 28 derived from euploid (EU79) and DS (DS18) lines. Data are presented as minimum to maximum, with notches centered on the median. ****p < 0.0001 via one-way ANOVA. Below heatmap expression of marker genes related to cilia. Values are shown as z-score. (**G**) Analysis of immunostained sections of euploids (EU79 and G22) and DS (DS18 and G21) CPCOs at day 56. Organoids are stained with the primary cilium marker ARL13B (Red) and the CP marker TTR (Green). The section was counterstained with Hoechst 33342 (Grey). Scale bar = 20 µm. Dotted white boxes denote areas that are magnified. (**H**) Stacked bar graph demonstrating the percentage of cells with a single cilium and multiple cilia in human CP cells in CPCOs derived from euploids (EU79 and G22) and DS (DS18 and G21) lines at day 56 of differentiation. Data are presented as mean ± standard deviation. Number of independent experiments = 3. *p < 0.05; **p < 0.01 via one-way ANOVA. (**I**) The distribution of cilia length in CPCOs derived from euploids (EU79 and G22) and DS (DS18 and G21) lines cultured for day 56. The data is presented as a percentage of CP cells with a single primary cilium or multi-cilia. *p < 0.01 via one-way ANOVA. (**J**) Immunostaining of euploid (EU79 and G22) and DS (DS18 and G21) CPCOs at day 56 for β-catenin (Red) and ZO1 (Green) and TTR (Magenta). All sections were counterstained with Hoechst 33342 (Blue). Scale bar = 25 µm. Dotted white boxes denote areas that are magnified and where the two fluorescent channels. Scale bar = 10 µm. (**K**) Bar graph showing the percentage of mislocalized ZO1 in euploid (EU79 and G22) and DS (DS18 and G21) CPCOs at day 56. Data are presented as mean ± standard deviation. ****p < 0.0001 indicates statistical significance via Student’s t-test. Number of independent experiments = 3.

We further discovered that relative to the euploid organoids, DS organoids showed significant up-regulation of genes associated with inhibitory neurons (e.g., *GAD1*, *ABAT*) at the expense of neural progenitors (e.g., *FOXP2*, *BCL11B*, *POU3F2*, *NES, NOTCH1*) and excitatory neuronal marker genes (e.g., *TUBB3*, *TH*, *SLC17A6*) (Figure S6G). To assess whether this influenced neuronal function, we again performed MEA analyses, revealing a significantly higher MFR in euploid organoids compared to DS (Figures S6H, S6I). We next used bicuculline, glutamate, and NMDA to evaluate responses from excitatory and inhibitory neuronal populations. The treatment with bicuculline significantly increased the firing rate in DS and euploid organoids compared to the baseline, and restored the electrophysiological imbalance in DS as compared to the euploid organoids (Figure S6J). In contrast, DS organoids treated with NMDA exhibited significantly less firing rate (∼ 1-fold) compared to the euploid (Figure S6J), in agreement with the under-representation of excitatory neurons in DS organoids (Figure S6G). Unlike NMDA, glutamate treatment showed elicited degrees of firing rate responses between the two groups (Figure S6J). These data therefore demonstrated that DS CPCOs recapitulate similar neural network hypo-excitability phenotypes as those observed in DS mouse models ^49^. We further noted that all marker genes of oligodendrocyte precursor cells (OPCs) were downregulated in DS organoids at 28 days (Figure S6B), and this was corroborated at the protein level via western blotting (Figure S6C). Immunofluorescence staining of organoid sections identified clear expression of SOX10 protein in cells within the cerebral cortex domain of euploid CPCOs (Figure S6D), and quantification demonstrated a significant reduction in SOX10 expressing cells in DS CPCOs as compared to their euploid counterparts (Figure S6E). This prompted us to examine the expression of the HSA21 gene *OLIG2* that plays a role in oligodendrogenesis in the developing dorsal forebrain ^50^, but found that this gene was under-expressed at the mRNA level in DS CPCOs (Figure S5D). Western blot and immunofluorescence staining of organoid sections next confirmed reduced protein expression and a reduced number of cells expressing OLIG2 in DS CPCOs as compared to euploid organoids, respectively (Figures S5E&F). In addition, we noted dysregulated expression of genes associated with interneuron specification in DS CPCOs as compared to euploid organoids (Figure S5G), consistent with previous findings ^20, 51, 52^. Collectively, these findings indicate that DS CPCOs recapitulate key aspects of DS brain pathology.

### SARS-CoV-2 productively infects Down syndrome CPCOs

Given that DS individuals are more susceptible to SARS-CoV-2 ^23, 24^, we next wished to assess the utility of CPCOs for modeling of SARS-CoV-2 infection of the human CNS. We first interrogated our RNA-Seq datasets of uninfected euploid and DS CPCOs for expression of the host genes that determine SARS-CoV-2 cell entry. SARS-CoV-2 can penetrate susceptible cells via the endosomal entry pathway, which requires the virus binding receptor angiotensin converting enzyme-2 (ACE2), and is facilitated by cleavage of viral Spike protein by the protease Furin. Alternatively, the virus can enter the cells via the cell surface pathway (direct fusion), which additionally requires cleavage of the spike protein by transmembrane serine protease 2 (TMPRSS2) ^53^. Our RNA sequencing data revealed robust expression of the *ACE2* gene in DS organoids at a level similar to that observed in euploid organoids (Figure 5A). As expected, for a gene located on HSA21, *TMPRSS2* expression was significantly higher in DS than in euploid organoids (Figure 5A). We further found that *FURIN* expression in DS organoids was 3-fold increased as compared to the euploid organoids (Figure 5A) even though *FURIN* gene is not located on HSA21. Western blot analysis corroborated the higher levels of TMPRSS2 and FURIN abundance in DS organoids as compared to euploid organoids (Figure 5B). Immunostaining similarly confirmed much higher expression levels of ACE2, TMPRSS2, and FURIN in CP than in the cerebral cortex portion of the CPCOs organoids (Figures 5C, S6K). These data therefore indicated that our CPCOs express receptors and proteases required for SARS-CoV-2 infection, and thus would likely constitute a suitable experimental model for studying SARS-CoV-2 CNS infection in individuals with DS. It also suggests that CP may facilitate virus entry into CNS as it expressed higher levels of *TMPRSS2* and *FURIN* than neurons.

**Figure 5.**
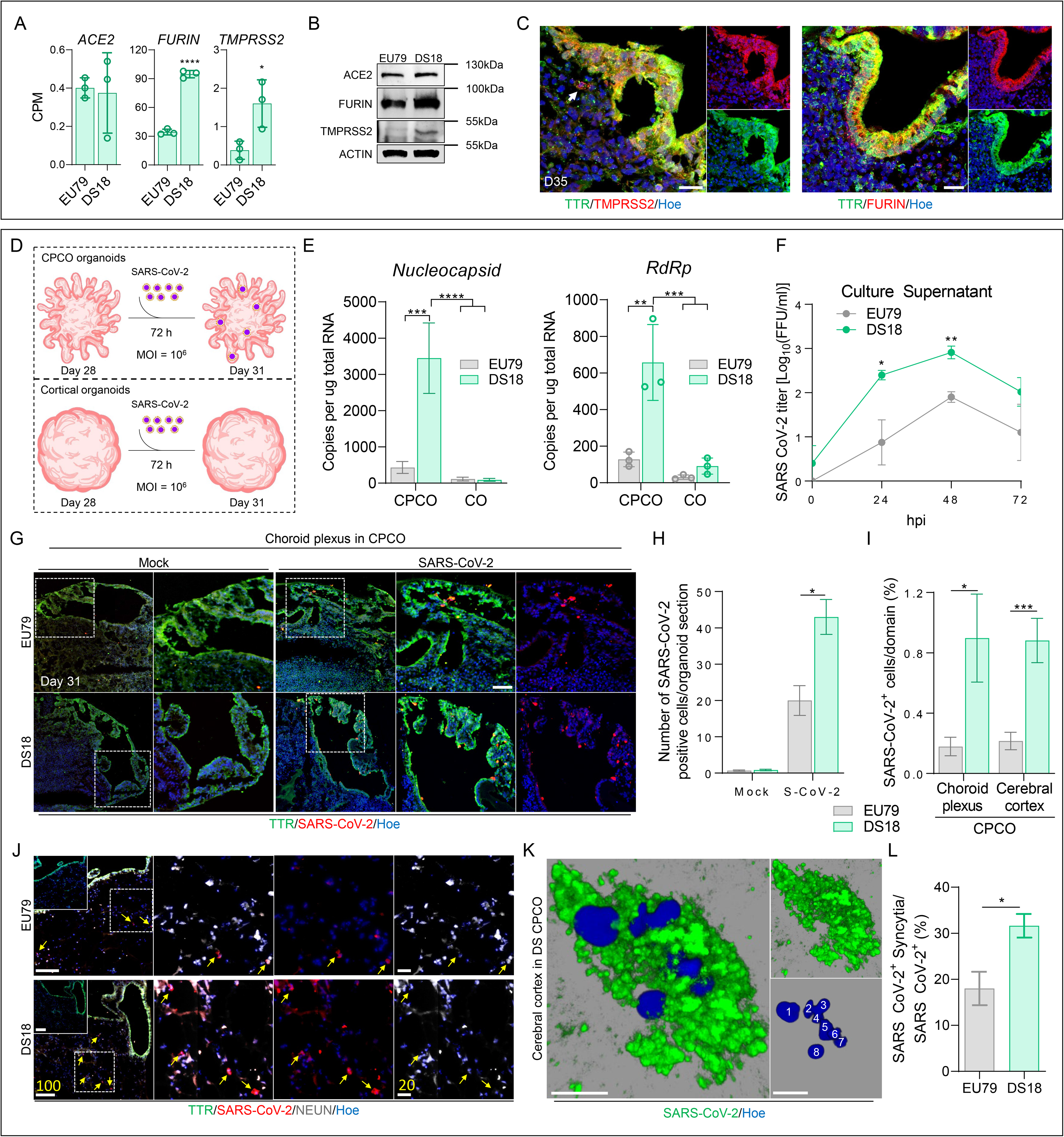
SARS-CoV-2 productively infects CPCOs but not traditional “naked” organoids. (**A**) Abundance of *ACE2*, *FURIN*, and *TMPRSS2*-derived RNA-Seq reads in euploid (EU79) and DS (DS18) CPCOs at day 28. Read counts were normalized to the library sizes. Data are presented as mean ± standard deviation. *p < 0.05, ****p < 0.0001 via Student’s t-test. Number of independent experiments = 3. (**B**) Western blots showing the protein levels of ACE2, FURIN, TMPRSS2, in euploid (EU79) and DS (DS18) CPCOs at day 31. Actin was used for normalization. All blots derived from the same experiment and processed in parallel. (**C**) Representative image of euploid-derived CPCOs at day 35 immunostained with TMPRSS2 (Red), FURIN (Red), and TTR (Green) antibodies. All sections were counterstained with Hoechst 33342 (Blue). Scale bar = 20 µm. (**D**) Schematic diagram of the SARS-CoV-2 infection protocol of CPCOs and traditional “naked” COs. (**E**) Bar graphs showing the amount of SARS-CoV-2 related genes *nucleocapsid* and *RdRp* in euploid (EU79) and DS (DS18) CPCOs and COs at day 31. Data are presented as mean ± standard deviation. **p < 0.01, ***p < 0.001, ****p < 0.0001 via 2-Way ANOVA. Number of independent experiments = 3. (**F**) Quantification of viral titers from euploid (EU79) and DS (DS18) CPCOs culture supernatants after SARS-CoV-2 (10^6^ FFUs) treatment of day 28 CPCOs at 0, 24, 48 and 72 hpi. Data are presented as mean ± SEM. N = 3 biological replicates consisting of 3 organoids each; *p < 0.05; **p < 0.01 via Student’s t-test. (**G**) Representative confocal images of day 31 CPCOs derived from euploid (EU79) and DS (DS18) immunostained with TTR (Green) and SARS-CoV-2 spike (Red) after SARS-CoV-2 (10^6^ FFUs) or mock (SARS-CoV-2 at 10^4^ FFUs). All sections were counterstained with Hoechst 33342 (Blue). Scale bar = 60 µm. (**H**) Bar graph showing the frequency of SARS-CoV-2-positive cells in CPCOs derived from euploid (EU79) and DS (DS18). Data are presented as mean ± SEM. *p < 0.05 via Student’s t-test. 15 CPCOs sections from three (n = 3) independent samples were examined. (**I**) Bar graph showing the percentages of SARS-CoV-2-positive cells within the choroid plexus and cerebral cortex domains of CPCOs derived from euploid (EU79) and DS (DS18). Data are presented as mean ± SEM. *p < 0.05, ***p < 0.001 via Student’s t-test. 15 CPCOs sections from three (n = 3) independent samples were examined. (**J**) Representative confocal images of day 31 CPCOs derived from euploid (EU79) and DS (DS18) immunostained with TTR (Green), SARS-CoV-2 spike (Red), and NEUN (Grey) after SARS-CoV-2 (10^6^ FFUs). All sections were counterstained with Hoechst 33342 (Blue). Scale bar = 100 µm. Scale bar for magnified images = 20 µm for scale bar. Yellow arrows indicate infected neurons. (**K**) High-resolution imaging and 3D reconstruction of SARS-CoV-2-positive syncytia with maximum Z-planes projection to highlight multiple nuclei. White numbers mark the individual nuclei. Scale bar = 10 µm. (**L**) Bar graphs showing the percentage of SARS-CoV-2-positive syncytia among total SARS CoV-2-positive cells after SARS-CoV-2 (10^6^ FFUs) treatment of day 28 CPCOs at 72 hpi. Data are presented as mean ± SEM. *p < 0.05 via Student’s t-test. 15 CPCOs sections from three (n = 3) independent samples were examined.

To assess the effect of CP on susceptibility of brain organoids to SARS-CoV-2 infection we infected day 28 euploid and DS CPCOs with 10^^6^FFU of SARS-CoV-2 for 72h (Figure 5D) and quantified the expression of SARS-CoV-2 *nucleocapsid*, *RdRp* (Figure 5E), *Envelope*, and *Spike* genes (Figure S6L) via qRT-PCR. This revealed that traditional “naked” COs were poorly infected by SARS-CoV-2 as compared to CPCOs and that DS CPCOs showed significantly higher expression of viral genes than euploid CPCOs. To confirm productive SARS-CoV-2 infection, we examined viral titers in the culture supernatants at 0, 24, 48, and 72 hours post infection (hpi), revealing a significant increase in titers of infectious virus in DS than in euploid group at 24 and 48 hpi (Figure 5F). Next, we examined the spatial distribution of SARS-CoV-2-infected cells in CPCOs. Upon SARS-CoV-2 infection, DS organoids showed a significantly higher proportion of cells expressing SARS-CoV-2 spike protein than euploid organoids (Figures 5G, H), consistent with higher viral RNA levels (Figures 5E, S6L). Given that CPCOs comprise both CP and cerebral cortex tissues, and since SARS-CoV-2 relevant receptor and proteases are more highly expressed in CP than in cortical tissues (Figures 5C, S6K), we next compared the cellular infection vulnerability to SARS-CoV-2 in CP versus cerebral cortex within CPCOs. Consistently, the number of SARS-CoV-2 spike protein positive cells in DS CP and cerebral cortex tissues was significantly higher than those in euploid organoids (Figure 5I). We interpreted these data to mean that infection of cerebral cortex cells is enabled by the presence of the CP population. We further stained these organoids with a pan-neuronal marker (NEUN) and found many cortical neurons in the cerebral cortex had been infected with SARS-CoV-2 (Figure 5J, yellow arrows). Interestingly, we observed that many of these infected neurons exhibited fragmented nuclei (Figures S7D&E), suggesting that infection with SARS-CoV-2 induces neuronal cell death.

Next, we examined the cellular consequences of SARS-CoV-2 infection of CPCOs. We observed the presence of syncytia in SARS-CoV-2-infected cells (Figure 5K), which was significantly increased in DS compared to the euploid CPCOs (Figure 5L). Interestingly, the majority of syncytia were observed in the cerebral cortex infected cells of DS (Figure 5K). By examining individual confocal Z-planes, we could identify as many as 8 nuclei within single infected DS cortical cells at 72 hpi (Figure 5K). Collectively, our data show that by using CPCOs as a 3D human cellular model, we were able to reveal significant SARS-CoV-2 tropism for CP cells that results in productive infection of cerebral cortex cells, and increased syncytia in DS that may promote viral spread through cell-cell fusion.

### Transcriptional Dysregulation of Down syndrome CPCOs upon SARS-CoV2 infection

To gain additional insight into the cellular responses to SARS-CoV-2 infection in CPCOs we performed RNA-seq of day 31 euploid and trisomy 21 CPCOs that were exposed to SARS CoV-2 for 72 hours. We first examined the reads that map to the SARS-CoV-2 genome and found a significantly higher load of SARS-CoV-2 RNA in DS CPCOs as compared to the euploid organoids (Figure 6A). The majority of virus-derived reads mapped to the 3’terminal part of the viral genome, which represents a subset of actively transcribed subgenomic RNAs, thus further confirming viral RNA replication in the infected cells (Figure 6B). Principle component analysis showed good clustering of biological replicates within different groups and separation of samples based on genotype in the first dimension and SARS-CoV-2 infection along the second dimension (Figure 6C). This analysis further revealed that the euploid and trisomy 21 SARS-CoV-2 positive organoids, were transcriptionally different from uninfected organoids in both groups (Figure 6C). Hierarchical clustering analysis using Spearman rank correlation of the top 500 most variable genes demonstrated that the SARS-CoV-2 positive organoids in the euploid and DS groups were transcriptionally different from the other two SARS-CoV-2 negative groups (Figure S7A). Compared to the euploid organoids with SARS CoV-2, we also confirmed the significant up-regulation of *TMPRSS2* and *FURIN* (Figure S7B), while *ACE2* remained at a comparable expression level between the two groups (Figure S7B). Comparison of SARS-CoV-2-infected and control uninfected DS CPCOs revealed 1058 upregulated genes and 215 downregulated genes (Figure 6D, Table S2). Comparison of SARS CoV-2-infected and control uninfected euploid CPCOs revealed 1118 upregulated genes and 758 downregulated genes (Figure 6E, Table S3), indicating that SARS-CoV-2 infection leads to large-scale transcriptional dysregulation.

**Figure 6.**
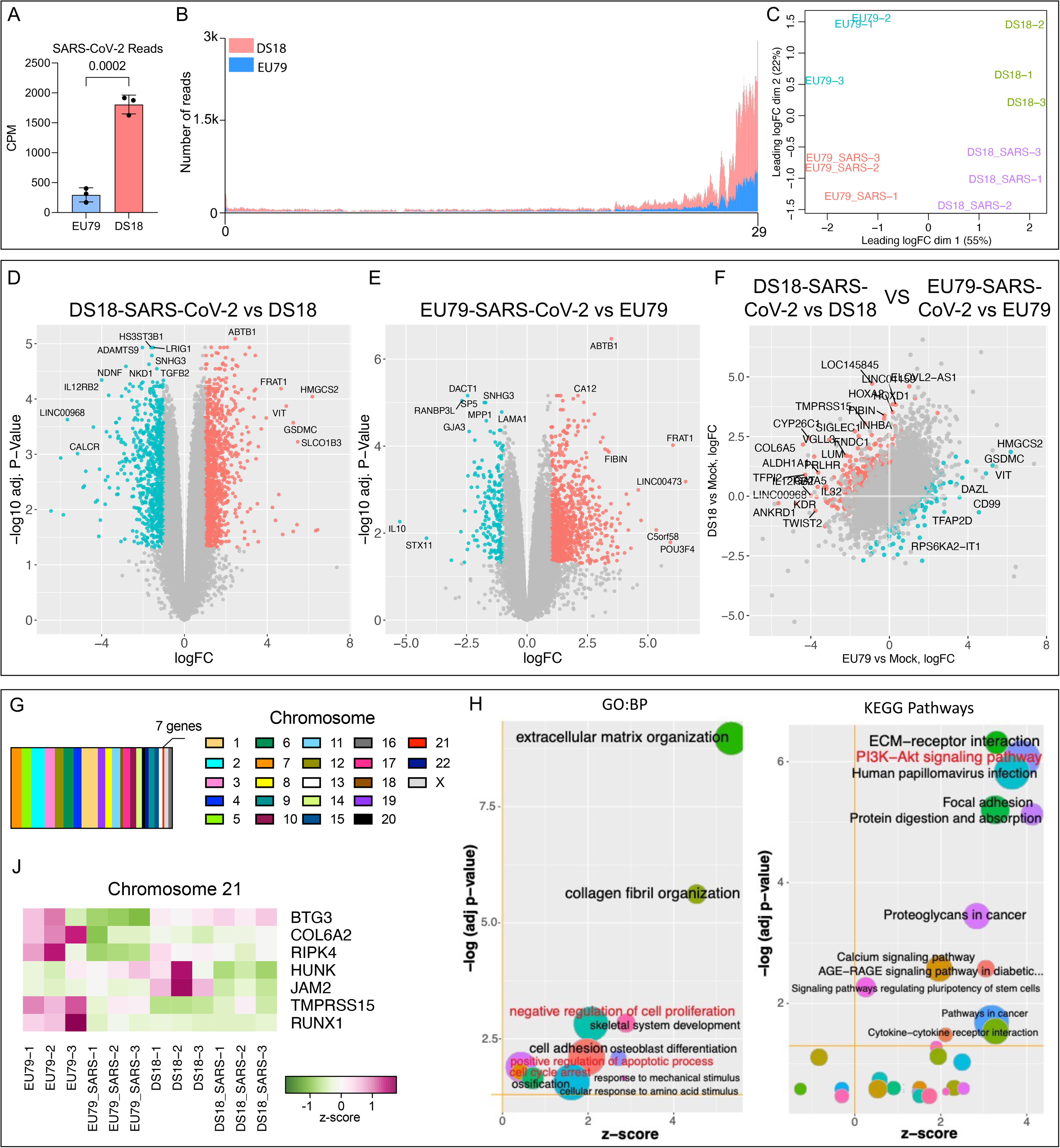
Transcriptional Analysis in Euploid and Trisomy 21 CPCOs upon SARS-CoV-2 infection. (**A**) Abundance of SARS-CoV-2-derived RNA-Seq reads in euploid (EU79) and DS (DS18) CPCOs. Read counts were normalized to the library sizes. Error bars indicate for SD. Statistical analysis was performed by Student’s t-test. (**B**) Distribution of SARS-CoV-2-derived RNA-Seq reads across viral genome. Data for DS (DS18) CPCOs is shown in red, and euploid (EU79) CPCOs is in blue. (**C**) Distribution multidimensional scaling analysis of RNA-Seq read counts derived from uninfected and SARS-CoV-2-infecetd human CPCOs of euploid (EU79) and DS (DS18). (**D, E**) Volcano plots highlighting differentially expressed genes (DEGs) in SARS-CoV-2 infected DS (**D**) and euploid **(E)** CPCOs compared to mock. (**F**) Scatter plot indicating the differences in SARS-CoV-2-induced gene expression changes between DS18 and EU79 CPCOs. In (D-F) the significantly (FDR-adjusted P-value < 0.05) up- and downregulated genes with at least 2-fold change in expression levels are shown in red and cyan, respectively. Most differentially expressed genes are labelled. (**G**) Distribution of the SARS-CoV-2-responsive differentially expressed genes identified in (F) across individual chromosomes. (**J**) Expression of the chromosome 21 – associated genes identified in (F) in SARS-CoV-2 infecetd and uninfected DS18 and euploid (EU79) CPCOs. (**H**) Gene ontology (GO) and KEGG pathway enrichment analysis of differentially expressed genes in (**F**). Z-scores indicate the cumulative increase or decrease in expression of the genes associated with each term. Size of the bubbles is proportional to the number of DEGs associated with respective GO term. BP – biological processes.

We next investigated unique SARS-CoV-2 responsive genes that act in DS but not in euploid organoids. We identified 392 upregulated genes and 219 downregulated genes (Figure 6F, Table S4). Notably, these genes were distributed across the entire genome with only seven of these located on chromosome 21 (Figures 6G, 6J). The gene ontology and pathway enrichment analyses of the identified differentially expressed genes revealed that DS18 CPCOs exhibit significantly stronger activation of the processes related to negative regulation of cell proliferation, positive regulation of apoptotic process, cell cycle arrest and P13K-Akt signaling pathways (Figures 6H, S7C). Notably, no activation of interferon (IFN) pathway was evident in SARS-CoV-2-infected organoids and no difference in expression of IFN-stimulated genes (ISGs) was observed (Figure 6H, Table S4). This indicates that higher susceptibility of DS CPCOs to SARS-CoV-2 infection is unlikely to be caused by imbalance in IFN response associated with extra copies of IFN response genes present on HSA21.

Considering the lack of active IFN response and activation of pro-apoptotic pathways in the infected organoids (Figure 6H, Table S4), the rapid decline in SARS-CoV-2 viral titers at 72 hpi (Figure 5F) is most likely explained by rapid elimination of permissive cells through programmed cells death rather than by virus clearance via innate immune mechanisms. To address this, we further examined the expression of pro-apoptotic genes (Table S5) and assessed the rate of cell death before and after SARS-CoV-2 infection. Before SARS-CoV-2 infection, DS organoids showed low expression of genes associated with apoptotic processes (Figure S8A), consistent with previous report suggesting that DS cells are less sensitive to apoptosis ^54^. However, exposure of DS organoids to SARS-CoV-2 significantly altered the expression of the pro-apoptotic genes (Figure S8A). Immunostaining further demonstrated a significant increase in cleaved caspase-3 positive cells following SARS-CoV-2 infection in DS organoids as compared to euploid organoids (Figure S8B). This is consistent with previous studies that demonstrated that alteration in apoptosis is associated with neurodegeneration in DS brain ^55^. Upregulation of many genes associated with cell adhesion and ECM remodeling (Figures 6F, 6H) suggests that infected CP cells in CPCOs could signal the CSF barrier to promote SARS-CoV-2 invasion ^56, 57^. Collectively, these transcriptome analyses are consistent with the notion that SARS-CoV-2 productively infects DS CPCOs and leads to SARS-CoV-2 invasion, defects in cell cycle and proliferation, and increased cell death of cortical neurons.

### TMPRSS2 inhibitors reduce SARS-CoV-2 infection of Down syndrome organoids

The transcriptional dysregulation in DS due to the triplication of HSA21 likely results in higher risk for more severe COVID-19, which may at least in part be due to the increased production of TMPRSS2 ^30^. Since we detected a 3-fold increase of TMPRSS2 in DS organoids (Figures 5A-B), we next evaluated the effects of the TMPRSS2 inhibitors Avoralstat, Camostat, and Nafamostat on productive SARS-CoV-2 infection (Figure 7A). DS CPCOs were pretreated with inhibitors for 6 h at an effective dose, and treatment continued for the next 72 h during SARS-CoV-2 infection, as suggested in previous studies ^58, 59^. We next compared the viral titers at 24 and 48 hpi to those produced in SARS-CoV-2 infected euploid and DS18 organoids not treated with inhibitors. All inhibitors significantly reduced viral titers in the supernatants of DS CPCOs at 24 and 48 hpi to a level comparable to that in SARS-CoV-2-infected euploid organoids (Figure 7B). Nafamostat exhibited the strongest inhibition of SARS-CoV-2 infection in DS organoids compared to Avoralstat or Camostat (Figure 7B), and this resulted in complete elimination of infectious virus from organoids at 48 hpi (Figure 7B). Notably, treatment with any of the three TMPRSS2 inhibitors completely abolished virus replication in euploid organoids (Figure S8C). Collectively these results indicate that TMPRSS2-mediated virus pathway (direct membrane fusion) is the predominant entry way for SARS-CoV-2 infection in brain tissue.

**Figure 7.**
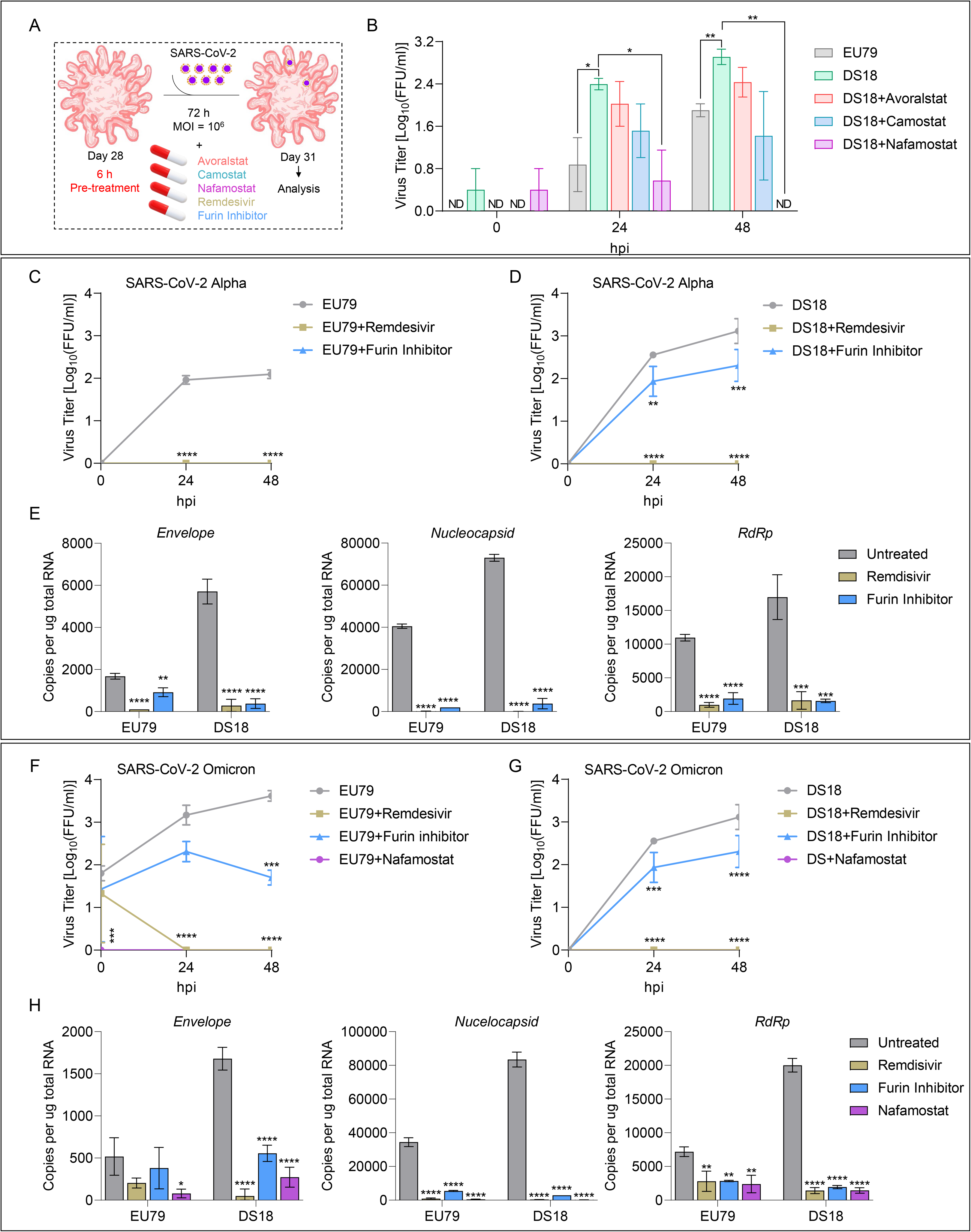
Drug Screening of FDA-approved Inhibitors against TMPRSS2 activity in CPCOs. (**A**) Schematic diagram presents the strategy of drug treatments and SARS-CoV-2 infection in CPCOs. TMPRSS2 and Furin inhibitors were added 6 hours before SARS-CoV-2 infection and for 72 hours during SARS-CoV-2 infection. Remdesivir was added immediately after SARS-CoV-2 infection and for 72 hours during SARS-CoV-2 infection. (**B**) Quantification of viral titers from CPCOs culture supernatants of euploid (EU79), DS (DS18), and DS18 treated with 100uM avoralstat or camostat or nafamostat after SARS-CoV-2 (10^6^ FFUs) infection of day 28 CPCOs at 0, 24, and 48 hpi. Data are presented as mean ± SEM. N = 3 biological replicates consisting of 3 organoids each; *p < 0.05; **p < 0.01 via Student’s t-test. (**C**) Quantification of viral titers from CPCOs culture supernatants of euploid (EU79) treated with Furin inhibitor (50 uM D-RVKR-CMK) and 100 100uM Remdesivir after SARS-CoV-2 (10^6^ FFUs) infection of day 28 CPCOs at 0, 24, and 48 hpi. Data are presented as mean ± SEM. N = 3 biological replicates consisting of 3 organoids each; ****p < 0.0001 via 2-Way ANOVA. (**D**) Quantification of viral titers from CPCOs culture supernatants of DS (DS18) treated with Furin inhibitor (50 uM D-RVKR-CMK) and 100 100uM Remdesivir after SARS-CoV-2 (10^6^ FFUs) infection of day 28 CPCOs at 0, 24, and 48 hpi. Data are presented as mean ± SEM. N = 3 biological replicates consisting of 3 organoids each; **p < 0.01; ***p < 0.0001; ****p < 0.0001 via 2-Way ANOVA. (**E**) Bar graphs showing the amount of SARS-CoV-2 related genes *Envelop*, *nucleocapsid* and *RdRp* in euploid (EU79) and DS (DS18) CPCOs at day 31. Data are presented as mean ± standard deviation. **p < 0.01, ****p < 0.0001 via One-Way ANOVA. Number of independent experiments = 3. (**F**) Quantification of viral titers from CPCOs culture supernatants of euploid (EU79) treated with Furin inhibitor (50 uM D-RVKR-CMK), 100 100uM Remdesivir and 100uM nafamostat after SARS-CoV-2 Omicron (10^6^ FFUs) infection of day 28 CPCOs at 0, 24, and 48 hpi. Data are presented as mean ± SEM. N = 3 biological replicates consisting of 3 organoids each; ***p < 0.0001; ****p < 0.0001 via 2-Way ANOVA. (**G**) Quantification of viral titers from CPCOs culture supernatants of DS (DS18) treated with Furin inhibitor (50 uM D-RVKR-CMK), 100 100uM Remdesivir and 100uM nafamostat after SARS-CoV-2 Omicron (10^6^ FFUs) infection of day 28 CPCOs at 0, 24, and 48 hpi. Data are presented as mean ± SEM. N = 3 biological replicates consisting of 3 organoids each; **p < 0.01; ***p < 0.0001; ****p < 0.0001 via 2-Way ANOVA. (**H**) Bar graphs showing the amount of SARS-CoV-2 Omicron related genes *Envelop*, *nucleocapsid* and *RdRp* in euploid (EU79) and DS (DS18) CPCOs at day 31. Data are presented as mean ± standard deviation. *p < 0.05, **p < 0.01, ****p < 0.0001 via One-Way ANOVA. Number of independent experiments = 3.

Considering that DS brain organoids also exhibited higher levels of Furin compared to euploid control (Figures 5A-D), we then assessed the effect of Furin inhibitor Decanoyl RVKR-CMK on viral replication in DS and euploid organoids. Furin cleavage of spike is not prerequisite for SARS-CoV-2 entry, but it highly facilitates viral infectivity ^60^. Treatment with Furin inhibitor completely abolished replication of SARS-CoV-2 in euploid organoids (Figure 7C&7E), which indicates that in brain tissue Furin activity is essential for SARS-CoV-2 entry via membrane fusion. The same effect was observed after treatment with FDA-approved COVID19 drug Remdesivir (Figure 7 C&7E). However, the effect of Furin inhibitor on virus replication in DS organoids was less profound. Although Decanoyl-RVKR-CMK caused a significant 10-fold reduction of viral titers at 24 and 48hpi, it did not eliminate the virus completely, while remdesivir did (Figure 7D-E). This can be explained by higher availability of TMPRSS2 and Furin in DS CPCOs (Figures 5A-C). On the other hand, the evolving SARS-CoV-2 omicron variants are less efficiently cleaved by Furin but exhibit greater sensitivity to TMPRSS2 inhibitor ^61^. This promoted us to assess the effect of Furin inhibitor on Omicron BA.5 replication in DS and euploid organoids. Both Nafamostat and Remdesivir treatments completely abolished Omicron BA.5 viral replication in DS and euploid organoids (Figures 7F-H). Although Furin inhibitor significantly reduced viral replication in both groups (Figures 7F&G), Furin inhibitor failed to completely abolish Omicron BA.5 in contrast to Nafamostat and Remdesivir (Figures 7F-H). Collectively, these experiments provide a proof-of-concept that CPCOs can be used as a model to screen for drugs that can inhibit SARS-CoV-2 neuropathology in specific genetic backgrounds, in this case for trisomy 21.

## DISCUSSION

Congenital disorders encompass a wide range of CP pathologies such as CP cysts, diffuse villous hyperplasia, lipoma and Sturge-Weber syndrome ^62^. For instance, CP cysts are common in fetuses with trisomy 18 and Aicardi’s syndrome, where CP cysts exhibit accumulation of the CSF causing hydrocephalus ^63^. Many of these CP abnormalities are linked to other structural anomalies in the brain, including cortical development. The development of human *in vitro* models that enable investigation of the cellular and molecular mechanisms underlying such diseases, particularly those in which the early stages of CP development and function are impaired, is therefore valuable, and may allow the development of therapeutic strategies that mitigate disruption of the CP function during development or later in life. In this study, we developed a rapid and robust protocol that generates organoids with multiple CPs that form ventricles that enclose developing functional cortical neuronal cells. We show that the CP epithelial layer that surrounds these organoids exhibits the typical features of the mature *in vivo* CP both at the cellular and molecular levels. The cortical cells (progenitors and neurons) and CP cells of these CPCOs arise from a common progenitor, the hNEct cells, that can be readily expanded and genetically manipulated with CRISPR to create various disease models.

Organoids representing multiple brain domains have been established ^64, 65^, including the CP ^9, 10^. Pellegrini and colleagues showed the presence of cortical neurons in their CP organoids using scRNA-seq ^9^ and CP organoids were subsequently utilized to demonstrate the susceptibility of CP to SARS-CoV-2 pathology ^10, 66^. BMP4 signaling is known to instruct neuroepithelia to become CP ^8^ at the expense of neural lineages ^9, 10^. Recent studies utilized a high concentration of BMPs to generate organoids that almost entirely consisted of CP cells ^9, 10^. In contrast, we used an approximately 10-fold lower BMP4 concentration to allow neural lineages specifications and further differentiation into cortical neural cells while still permitting CP development in CPCOs. Consistent with this notion (Figure S1D), we demonstrated that different concentrations of BMP4 altered the content of neural progenitors. Recently, the *in vitro* generation of hNEct from hPSCs has been instrumental in the production of forebrain cells in conventional monolayer culture and in 3D organoid models ^67–69^. Here, we took advantage of the developmental potential of hNEct sheets to generate a complex 3D CPCO model in which two different brain domains, cortical neurons and CP epithelium, develop in parallel from common hNEct progenitors that self-organize, and interact to promote the formation of ventricle-like structures. Different hPSC lines were used to generate CPCOs, demonstrating the reproducibility of this protocol. The CPCOs contain key components of the mature CP *in vivo* including the establishment of apicobasal polarity. We also detected high mitochondrial density in CPCO CP epithelium that *in vivo* provides the metabolic capacity for maintaining the secretory activities and ionic gradients across brain-CSF barriers ^70^. Transcriptomic data of CPCOs over time suggest the ability of CPCOs to produce CSF, and our multiplexed ELISA analyses of CST3, APP and B2M in CPCOs-medium confirm secretion of these CSF proteins. We also showed that the evolutionary conserved anti-aging protein KLOTHO is expressed in human CPs of CPCOs. Klotho is known to protect against multiple neurological and psychological disorders ^33^. We further showed that human CP cells in CPCOs projected up to 9 cilia per cell during differentiation, with a significant shift from primary to multi-cilia during differentiation. Whether these cilia are of the motile 9+2 axoneme kind or non-motile 9+0 axonemal cilia remains to be determined. The CPCOs reported here should provide a useful model to address the axoneme organization and diversity in subsets of human CP cells, and may illuminate their respective roles in CP function and cortical development.

Indeed, the presence of developing functional cortical cells that are juxtaposed to CP cells in CPCOs has the distinct advantage that it allows investigation of CP mediated neurodevelopmental defects such as ventriculomegaly in DS and hydrocephalus in Bardet Biedl syndrome through the use of patient specific iPSCs lines or genome edited control iPSC. Because such neurodevelopmental defects can occur very early during embryo development, they have been difficult to study in a human setting and our model offers unique opportunities to uncover novel disease processes and to assess the effectiveness of pharmacological interventions designed to rescue CP malformations and promote normal cortical brain development. The future challenge will be to control anterior-posterior identity of hNEct derived CP while maintaining maturation of cortical neurons. The ability to fuse different domains of the brain using the organoids system has been recently achieved and was used to analyze complex neurodevelopmental defects ^71, 72^. Similarly, the establishment of CPCOs with a lateral ventricle identity affords the potential to establish the fourth ventricle CPs with hippocampus, for example through judicious sonic hedgehog activation, and may offer an opportunity to establish and study the anterior-posterior identity with complex networks in an all-brain *in vitro* 3D model in the future.

CPCOs contain key, functional components of the forebrain tissue; they are highly reproducible and can be easily maintained in culture for several months. These attributes make CPCOs an attractive system to study neurodevelopmental defects. As a proof of principle, we have modeled DS, a prevalent genetic disorder that perturbs forebrain development, displays ventriculomegaly linked to defective cilia function and thus results in intellectual disability. Consistent with previous reports, CPCOs derived from trisomy 21 iPSCs displayed imbalanced production of excitatory and inhibitory neurons as compared to the euploid counterparts. Our MEA data in DS CPCOs are largely in agreement with the impaired neuronal function and hypo-excitability associated with DS, and drugs such as NMDA and bicuculline restored the electrophysiological imbalance, consistent with previous reports in animal models ^49, 73^. CPCOs would provide an attractive platform for assessing the effectiveness of different pharmacological agents aimed at stabilizing the electrophysiological imbalance in DS brain to enhance forebrain function.

In the developing forebrain, Olig2 is expressed ventrally in neural stem cells, and later in development is expressed dorsally to give rise to oligodendroglial cells ^50^. OLIG2 is a HSA21 gene that was recently shown to be overexpressed in neural stem cells populations of ventral forebrain organoids derived from DS iPSCs ^20^. In contrast, we rather found downregulation of OLIG2 in CPCOs derived from DS iPSCs at both the gene and protein levels as compared to euploid organoids. It is not unlikely that this difference is due to the dorsal forebrain identity of the CPCOs generated in this study and we hypothesize that the downregulation of OLIG2 in DS CPCOs may underlie downregulation of OPC markers (*SOX10* and *PDGFRA*) in this model. We further discovered defective cilia formation and disrupted CP polarity in CPCOs derived from the DS lines. Interestingly, we found that CPCOs derived from different DS iPSCs lines exhibited different penetrance degrees of DS pathology. This is consistent with previous studies indicating variable disease penetrance in individuals with DS ^74^, further supports the notion that the severity of DS phenotypes is affected by genetic background. Down syndrome foetuses and indeed 2D neuronal cultures and cortical brain organoids derived from DS hiPSC exhibit an increased production of astrocytes ^15, 75^. In contrast, analysis of our RNA-seq data of top 500 most variable genes did not detect changes in the expression level of genes associated with astrocytes (Table S1). We speculate that this inability to phenocopy this aspect of DS brain development with CPCOs may be due to the fact that the CPCOs protocol involves prolonged exposure to BMP4 (Figure 1A), which is known to promote astrocyte differentiation ^76^ and therefore may have obscured intrinsic differences in astrocytes production between DS and euploid groups. Defects in the development and function of cortical interneurons are linked to many disorders characterized by intellectual disability, and defective development of cortical interneurons was reported to be impaired in Down syndrome ^51^. Previous studies using human iPSC-based organoid models of Down syndrome^20^ and postmortem Down syndrome brain samples ^20, 51^ demonstrated deficits in calretinin interneurons ^20, 51, 52^, consistent with our findings that calretinin interneurons encoded gene *CALB2* was significantly downregulated in CPCOs derived from Down syndrome iPSCs (Figure S5G). We also found that Down syndrome CPCOs showed altered expression of interneurons subtypes-specific genes, with high expression of calbindin (*CALB1*), and downregulation of calretinin (*CALB2*) and Neuropeptide Y (*NPY*) genes (Figure S5G). In agreement with a previous study ^20^, our analysis also failed to detect significant differences in the parvalbumin interneurons expressed gene *PVALB*.

Viral infection acquired antenatally can have devastating impacts on the developing fetal brain ^77^. Several clinical reports of infants born to women with SARS-CoV-2 infection during pregnancy displayed developmental delay in 10% of infants at 12 months of age ^78, 79^. Notably, the neurodevelopmental morbidity was not associated with prematurity, suggesting a specific mechanisms of SARS-CoV-2 neurotropism rather than simply contributing to pregnancy complications ^80^. *In vitro*, SARS-CoV-2 was shown to have neuroinvasive and neurotropic attributes for the human CP ^10^, neurons ^81^, and astrocytes ^82^, indicating the potential of SARS-CoV-2 to impact early brain development. Given the fact that *TMPRSS2* is located on chromosome 21, and that epidemiological studies have found a distinct vulnerability of people with DS towards SARS-CoV-2, as illustrated by the 4-fold increased risk for COVID-19-related hospitalization and 10-fold increased risk of death ^23, 24^, we utilized our DS CPCOs as a model to study SARS-CoV-2 infection. Upon exposure to SARS-CoV-2, we observed robust infection of the CP in CPCOs-derived from DS iPSCs and consequent invasion into the cerebral cortex. Since the traditional “naked” COs were poorly infected, it is likely that CP cells in CPCOs serve as viral ‘replication hubs’ that support viral invasion and spread to other cortical cells, consistent with a previous study indicating that COVID-19 neurological symptoms in post-mortem brains are linked to perturbations in barrier cells of the choroid plexus ^83^. We confirmed that in this model TMPRSS2 and FURIN, but not ACE2 and NRP1, are significantly up-regulated in CP of DS CPCOs. As discussed above, DS CPCOs also display defective epithelial polarity of the CP. The tight junctions in CP epithelial polarity serve as a barrier to invaders and hamper virus endocytosis. Therefore, both increased dosage of TMPRSS2 as well as defective tight junctions in CP of DS organoids may combine to promote SARS-CoV-2 entry and subsequent neuropathology. Gene expression studies demonstrated genome wide transcriptome deregulation in DS individuals, DS iPS cells, and the DS Ts65D mouse model ^84^. Individual with DS develop severe complications such as excessive immune response during viral respiratory infections ^28^. It was recently demonstrated that type I interferon (IFN-I)-expression in the CP is age dependent and negatively affects brain function^85^. IFN-I expression profile was also found in human CP ^85^. These data identified chronic aging induced IFN-I signature at the CP, which is often associated with antiviral response ^85^. On the other hand, individuals with DS develop severe complications during viral infection due to impaired immune response ^54^. In the absence of any detectable infections, DS individuals exhibit chronic hyperactive IFN response ^86^, and elevation of many cytokines and chemokines levels known to act downstream of IFN signaling ^87^. This is in a part due to the fact that IFN receptors (IFNRs) are encoded on HSA21 ^86^. While we found that several HSA21 genes (*MX2, TMPRSS2, ADAMTS5,* and *RUNX1*) associated with viral infection are expressed in CPCOs, we failed to detect deregulation of interferon receptors and interferon signaling pathway molecules, and a general lack of expression of interferon associated genes in our day 31 old CPCO and in SARS-CoV-2 infected CPCOs. This observation is consistent with the ability of the virus to evade and inhibit IFN response ^88, 89^. Instead, we observed activation of apoptotic pathways and death of infected CPCO cells (Figures S8A-B).

We further showed that inhibition of TMPRSS2 using FDA-approved drugs (Avoralstat, Camostat, Nafamostat) strongly decreased SARS-CoV-2 replication in all organoids, indicating that TMPRSS2 is required for viral entry into the cells in EU and DS CPCOs. Consistent with previous reports ^90^, we found a higher potency of Nafamostat for inhibiting SARS-CoV-2 infection and replication as compared to Avoralstat and Camostat. Human neurons have been previously reported to express low levels of ACE2 and no TMPRSS2 ^91^. In this study, we found that CP exhibits high expression of TMPRSS2 and Furin, which further increases in Down syndrome (Figures 5 A-B). Our data show that TMPRSS2 inhibition strongly impacts virus infection and replication, further underlining the significance of this receptor for SARS-CoV-2 infection of CPCOs, and providing additional evidence that TMPRSS2-expressing CP cells act a primary reservoir for SARS-CoV-2 infection and spread. Inhibition of Furin also substantially reduced viral replication in DS CPCO, and in euploid CPCO completely eliminated the virus. Given that cleavage of viral spike protein by Furin is known to facilitate viral entry, while not being absolutely essential, these data indicate that DS CPCO likely have sufficient amount of TMPRSS2 to enable viral entry in absence of the furin activity, however, in euploid organoids that have less TMPRSS2, Furin may however be strictly required for SARS-CoV-2 infectivity. In addition, we showed that treatment with FDA approved COVID-19 drug Remdesivir results in complete elimination of the virus from both euploid and DP CPCOS. This is consistent with a high efficacy of the drug against SARS-CoV-2 ^92^.

In conclusion, our study underscores the relevance of CPCOs for modelling neurodevelopmental diseases such as DS and (corona)virus research, and signifies the importance of protease proteins (TMPRSS2, FURIN) as attractive therapeutic targets for inhibiting SARS-CoV-2 neurotropism. CPCOs should further prove useful for identifying and screening therapeutics for future emerging viruses and for modelling congenital disorders that involve choroid plexus dysfunction.

## MATERIAL AND METHODS

### Human embryonic stem cells culture and cortical organoids generation

Human embryonic stem cells H9 from (Wisconsin International Stem Cell Bank, WiCell Research Institute, WA09 cells), WTC iPSC (gift from Professor Bruce Conklin), G22, DS18 and EU79 iPSCs lines (available in our lab) were cultured according to Stem Cell Technologies protocols that can be found in (https://www.stemcell.com/maintenance-of-human-pluripotent-stem-cells-in-mtesr1.html) on feeder free in hESCs medium on Matrigel (StemCell Technologies, Cat. #354277) in mTeSR (Stem Cell Technologies, Cat. #85851).

### Choroid Plexus-Cortical organoids generation

To generate CPCOs, hPSC colonies were plated on a hESC qualified basement membrane matrix (StemCell Technologies, Cat. #354277) in a 6-well plate at 20%-30% density. hPSC colonies were maintained with mTeSR for one day prior neuroectoderm (NEct) induction. To generate NEct colonies, hPSC colonies were cultured for three days in N2 medium: DMEM/F12 (Gibco, Cat. #11320-33), 2% B-27 supplement (Gibco, Cat. # 17504044), 1% N-2 supplement (Gibco, Cat. #17502-048), 1% MEM Non-Essential Amino Acids (Gibco, Cat. #11140-050), 1% penicillin/streptomycin (Gibco, Cat. #15140148), and 0.1% β- mercaptoethanol (Gibco, Cat. #21985-023), containing dual SMAD inhibitors, SB-431542 (10 μM) and LDN 193189 (100 nM). Fresh N2 medium with inhibitors were added daily. On the fourth day, induced NEct colonies were lifted using dispase (2.4 unit/ml) to form neural spheroids as we recently reported ^68^. For the next four days, these spheroids were cultured in N2 medium supplemented daily with (bFGF, 40 ng/mL; R&D, Cat. #233-FB-01M), 2µM CHIR99021 (Sigma Aldrich, Cat. # SML1046-5MG) and 2ng/ml of BMP4 (Thermofisher, Cat. # PHC9391). Patterned neural spheroids were then embedded in Matrigel (StemCell Technologies, Cat. #354277) and switched to the terminal differentiation medium DMEM-F12 (Gibco, Cat. #11320-33): Neurobasal medium (Gibco, Cat. #A35829-01) 0.5% N2 (Gibco, Cat. #17502-048), 12.5 µl of insulin in 50 ml media (Sigma), 1% GlutaMAX, 1% MEM Non-Essential Amino Acids (Gibco, Cat. #11140-050), 1% penicillin/streptomycin (Gibco, Cat. #15140148), 17.5 µl of β-mercaptoethanol in 50 ml media (Gibco, Cat. #21985-023), and 1% B-27 supplement (Gibco, Cat. # 17504044), supplemented with 3µM CHIR99021 and 5ng/ml of BMP4. Fresh media was replaced three times a week. All experiments were carried out in accordance with the ethical guidelines of the University of Queensland and with the approval by the University of Queensland Human Research Ethics Committee (Approval number-2019000159).

### qRT-PCR

Total RNA was isolated from organoids as described previously ^93^. For qPCR, 1µg of isolated RNA was utilized to generate the complementary DNA (cDNA) using the first strand cDNA Synthesis Kit (Thermo scientific, Cat. #K1612). SYBR Green (Applied Biosystem, Cat. #A25742) was utilized, and PCR standard reaction conditions were set according to the manufacturer’s instructions. For quantification of viral RNA 1ug of total RNA was reverse transcribed using qScript cDNA SuperMix (Quanta Bio, Cat. #95548). The cDNA was diluted 1:10 and 3 ul of the solution were used as a templated for qRT-PCR. The qPCR was performed using QuantiNova SYBR Green PCR Kit (Qiagen, Cat. # 208056) on Applied Biosystem QuantStudio 6 instrument. RNA copy numbers were determined by comparing sample Ct values to the standard curve obtained by amplification of the DNA standards generated by end point RT-PCR with the same sets of primers. PCR primers were designed using the NCBI free online system, all the RT-qPCR primers are listed in Table S6. All experiments were performed in biological triplicates for every sample, and the expression values were normalized against the GAPDH expression value of each sample. The means and standard deviations were calculated and plotted using the GraphPad Prism 9^®^.

### Immunohistochemistry

Tissue processing was performed as described in and immunohistochemistry (IHC) was performed as described in ^94^. In brief, organoids were fixed in 4% PFA for 60 min at RT, followed by washing with 1x phosphate buffer saline (PBS) three times for 10 min at RT. Fixed organoids were then immersed in 30% sucrose in PBS at 4 °C and allowed to sink before being embedded in a solution containing at 3:2 ratio of Optimal Cutting Temperature (O.C.T) and 30% sucrose on dry ice. Mounted tissues were then subjected to serial sections at 14-µM thickness and collected onto Superfrost slides (Thermo Scientific, cat. #SF41296). To performed IHC, sectioned organoids were washed three times with 1x PBS for 10 minutes at RT before blocking for 1 hour with 3% bovine serum albumin (BSA) (Sigma, Cat. A9418-50G) and 0.1% triton X-100 in 1x PBS. Primary antibodies were added overnight at 4 °C before washing three times with PBS for 10 minutes each at RT. For immunocytochemistry (ICC) ^95^, cells were allowed to be fixed with 4% PFA in 1x PBS for 10 min at RT. The cells were then washed three times with 1x PBS at RT before blocking and adding primary antibody as stated above. Tissues and cells were then incubated with appropriate secondary antibodies for one hour at RT before mounting and imaging. The wholemount was performed as described before^96^. All samples were counterstained with Hoechst 33342 (Invitrogen, Cat. #H3570). All images were acquired using confocal microscopy (Leica TCS SP8) based in SBMS Imaging Facilities based at the University of Queensland. The primary antibodies used in this study are listed in Table S7. Alexa-488, Alexa-546, and Alexa-633-conjugated secondary antibodies were obtained from Jackson ImmunoResearch Laboratory.

### RNA-sequencing

RNA was isolated from the pools of three individual organoids using TRIreagent (Sigma, USA) as described previously (Slonchak et al., biorxiv, 2021). RNA integrity was analyzed on TapeStation 4200 (Agilent, USA) and samples with RIN>8 were used for library preparation with TrueSeq RNA Library Preparation Kit v2, Set A (Illumina, USA). Barcoded cDNA libraries were pooled and sequenced on Illumina NextSeq 500 instrument using NextSeq 500/550 High Output 75 Cycles Kit v2.5 (Illumina, USA). Image acquisition, processing and fastq demultiplexing were performed using in-instrument software as described previously ^97^.

### Differential Gene Expression Analysis

Quality control of raw sequencing data was performed using FastQC software v.0.72. Reads were then trimmed using Trimmomatic software v.0.36.6 with the following parameters: ILLUMINACLIP : TruSeq3 - SE : 2 : 30 : 10, LEADING : 32 TRAILING : 32, SLIDINGWINDOW : 4 : 20, MINLEN : 16. Trimmed reads were mapped to SARS-CoV-2 genome (GISAID Accession ID: EPI_ISL_944644) using Bowtie2 v.2.4.4. Aligned reads were visualised and quantified using Integrative Genomic Viewer v.2.13.1 (Broad Institute, USA). Human reads were mapped to the genome assembly hg38 using HISAT2 v.2.2.1. Feature counting was performed using featureCounts v2.0.1 with counting mode set to “Union”, strand to “Unstranded”, feature type was “exon”, and ID attribute was Gene_ID.

Differential gene expression analysis was performed using edgeR v.4.2. Low abundance reads (<1 cpm) were removed from the data set and data was normalised to library sizes and composition bias using the trimmed mean of M-values (TMM) method. Normalised data was analysed by multi-dimensional scaling analysis and used to build quasi-likelihood negative binomial generalized log-linear model. The Treat test (glmTreat) was then applied to the contrasts CP-NC, DS18_CP-EU79_CP, SARS_DS18-Mock_DS18 and SARS_EU79-Mock_EU79 to identify the genes that are differentially expressed between the groups by at least one log2. The quasi-likelihood F-test (glmQLFTest) was applied to the contrast (SARS_DS18-Mock_DS18)-(SARS_EU79-Mock_EU79) to identify the genes affected by infection specifically in DS18 organoids. Genes were considered differentially expressed if FDR-corrected P-values were <0.05. Gene expression data were plotted using ggplot2 v.3.3.2. Gene ontology and KEGG pathway enrichment analyses were performed using Database for Annotation, Visualization and Integrated Discovery (DAVID) v6.8. Enrichment data was then combined with expression values and z-scores were calculated using the GOplot v.1.0.2 R package and plotted using ggplot2 v.3.3.2. Heat maps were generated using heatmap.2 function of R-package gplots v3.1.2.

### Transmission Electron Microscopy

Transmission electron microscopy were proceed according to ^98^. In brief, organoids were in 0.1 M Sodium cacodylate buffer in ddH20 containing 2.5% Glutaraldehyde and 2% Paraformaldehyde over night at 4°C. Organoids were first washed three times in 0.1 M Sodium cacodylate buffer for 10 min at RT, followed by immersing in 2% osmium tetroxide (2 ml 4% osmium tetroxide and 2 ml 0.2M cacodylate buffer) for 90 min at RT. The staining buffer was then replaced with 2.5% potassium ferricyanide (4.8 ml 0.2M cacodylate buffer and 4 ml 6% potassium ferricyanide) for 90 min at RT. The organoids were then washed in water three to five times at RT until the water is completely clear, before dipping the organoids in the thiocarbohydrazide solution (0.1g thiocarbohydrazide (locked cupboard) in 10 mL water) for 45 min at 40°C. To remove the background stain, organoids were washed four to five times in water at RT before immersing the organoids into 2% osmium tetroxide in 0.1M cacodylate buffer for 30 min at RT. Organoids were again washed in water to remove the black background. Washed organoids were then dipped into 1% uranyl acetate overnight at 4°C, followed by incubation in 0.03M aspartic acid solution for 30 min at 60°C. Incubated organoids were then washed three times for 30 min at RT. For embedding, the organoids were dehydrated in a series of ethanol 20%, 50%, 60%, 70%, 80%, 90% and 100% for 30 time at RT each. Organoids were then infiltrate with Durcupan resin for 12h at RT. Embedded organoids were then incubated for 48h in 60°C before trimming and imaging.

### Multiplexed ELISA

Organoids media for investigations of CSF biomarkers (CST3, APP, and B2M) levels were collected at days 7, 14, 21, 28, 42, and 56. The level of CST3, APP, and B2M in collected media were examined using Human Magnetic Luminex^®^ Assays (R&D Systems; Cat. # LXSAHM-03) according to the manufacturer’s instructions and read on a MAGPIX instrument (Luminex Corp).

### Multielectrode Array (MEA) Recording and Analysis

Whole organoids cultured in terminal differentiation medium were transferred prior to recording to the MEA plate (4096 channels at cellular resolution (20 µm) over a large area (up to 5.1 x 5.1 mm^2^), with a sampling frequency of 18kHz/ channel, 3Brain) and recorded in a BioCAM X system (3Brain). Media was removed from each well until only a thin layer remained to allow the attachment of organoids to the electrode grid under the 37 °C temperature and 5% CO2. During the recording, 50 µM Glutamate (Sigma Cat. #G1251-100G) or 10 µM NMDA (Sigma Cat. #M3262-25MG) were added to warm media to stimulate neuron activity in organoids. To record the inhibitory pharmacologic effects on CPCOs, 50 µM Bicuculline (Sigma Cat. #14340-25MG) was added to the media. The baseline activity was recorded without pharmacologic effects. Data was exported as .csv files for analysis using BrainWave Software. For the mean firing rate, the algorithm PTSD (Precise Timing Spike Detection) were applied having a threshold 8 times the standard deviation above the background noise with a 2 ms peak lifetime period and 2 ms refractory period were imposed after each detected spike. The bursts were detected using 2 ms of max spike interval and 5 as a minimum number of spikes.

### Western Blot

Western blotting was performed as described previously ^33^. In brief, Pierce^TM^ RIPA Buffer (ThermoFisher Scientific, Cat. #89900) containing a cocktail of protease and phosphatase inhibitors (Roche)) was used to lyse the organoids. A sonicator was used to sonicate the organoids, and Pierce^TM^ bicinchoninic acid (BCA) protein assay kit (ThermoFisher Scientific, Cat. #23227) was used to quantify protein concentration according to the manufacturer’s instructions. The extracted proteins were then heated for 10 min at 100°C before loading. Equal proteins amount was loaded and separated using Mini-PROTEAN TGX Stain-Free-Gels (BIO-RAD, Cat. #4568044). iBlot 2 PVDF Mini Stacks (Invitrogen, Cat. #IB24002) was used to transfer the separated proteins. 5% Skim Milk in TBS-T (20 mM Tris-HCl, pH 7.6, 136 mM NaCl, and 0.1% Tween-20) was used to block the membranes for 1 hour at RT, before incubation with primary antibodies which were diluted in 5% BSA for 12 hours at 4°C. The primary antibodies used in this experiment are listed in Table S6. Incubated membranes were then washed three times with 1x TBST for 10 minutes each at RT followed by another incubation with secondary antibody diluted 1:5000 in 5% Skim Milk in 1x TBST for 1 hour at RT. Finally, the incubated membranes were washed again three times with 1x TBST for 10 minutes each at RT and visualised with Clarity Western ECL Substrate (BIO-RAD, Cat. #170-5060).

### SARS-CoV-2 infection of CPCOs

SARS-CoV-2 isolate QLD1517/2021 (alpha variant, GISAID accession EPI_ISL_944644), passage 2, was recovered from nasopharyngeal aspirates of an infected individual and provided by the Queensland Health Forensic and Scientific Services, Queensland Department of Health. The obtained isolate was amplified on Vero E6-TMPRSS2 cells to generate virus stock. Virus titers were determined by immuno-fluorescent foci-forming assay on Vero E6 cells. Organoids were incubated in 500 μl media containing 10^6^ foci-forming units (FFU) of the virus for 6 hours at 37 °C, then inoculum was removed, organoids were washed 3 times and then maintained in 1 ml of fresh medium. Culture fluids were sampled at 0, 24, 48 and 72 hpi to monitor virus replication. Organoids were pre-incubated for 6 h in media containing 100uM TMPRSS2 inhibitor (Avoralstat, Camostat or Nafamostat) or 50 uM D-RVKR-CMK. After drug treatment organoids were infected overnight with SARS-CoV-2 at the dose of 10^6^ foci-forming units (FFU) per organoid. 100uM Remdesivir was added immediately after SARS-CoV-2 infection. Inoculum was then removed, organoids were washed three times and placed into the media containing the same concentration of corresponding drug.

### Immuno-fluorescent foci-forming assay

Ten-fold serial dilutions of cell culture fluids were prepared in DMEM media supplemented with 2% FBS and 25 µl of each dilution were used to infect 2x10^4^ Vero E6 cells pre-seeded in 96 well plates. After 1 h of incubation at 37 °C with the inoculum, 175 µl overlay media was added to each well. The overlay media consisted of 1:1 mixture of M199 medium (supplemented with 5% FCS, 100 μg/ml streptomycin, 100 U/ml penicillin, and 2.2 g/L NaHCO3) and 2% carboxymethyl cellulose (Sigma-Aldrich, USA). At 1 day post-infection, overlay medium was removed and cells were fixed by submerging in cold 80% acetone (diluted in 1x PBS) for 1h at -20°C. The following procedures were handled in PC2. The cell monolayer was dried and blocked for 60 min with 150 µl/well of Clear Milk Blocking Solution (Pierce, USA). Viral spike protein was probed by incubation with 50 μl/well of CR3022 mouse monoclonal antibody diluted 1:1000 for 1h, followed by five washes with phosphate buffered saline containing 0.05% Tween 20 (PBST), 1h incubation with 50 μl/well of 1:1000 dilution of goat anti-mouse IRDye 800CW secondary antibody (LI-COR, USA) and another five PBS-T washes. All antibodies were diluted with Clear Milk blocking buffer (Pierce, USA) and incubations were performed at 37 °C. Plates were then scanned using an Odyssey CLx Imaging System (LI-COR) using the following settings: channel = 800 and 700, intensity = auto, resolution= 42 μm, quality= medium and focus= 3.0 mm. Virus replication foci were then counted, and viral titers were calculated.

### Statistical analysis

Normally distributed data were expressed as the mean ± standard deviation of the mean of independent experiments. For non-normally distributed data, the median ± standard deviation was used to express the values. The number of biological replicates as well as the sample size are indicated in the figure legends. The Student’s t-test and one-way or two-way ANOVA were utilized for comparing two and more than two groups, respectively. The Tukey’s post-hoc analysis was applied for comparisons to a single control. Statistical analysis was performed using GraphPad Prism 9^®^ software. Minimal statistical significance was defined at P < 0.05.

## ACKNOWLEDGMENTS

M.R.S. is supported by the Children Hospital Foundation (PCC0252021). E.J.W. is supported by the MRFF Leukodystrophy Flagship – Massimo’s Mission (EPCD000034) and by the UQ Centre for Stem Cell Engineering and Regenerative Engineering (UQ StemCARE). A.A.K. is supported by NHMRC Ideas Grant (2012883) and Australian Infectious Diseases Research Center (AID) Seed Grant. This work was funded by UQ Centre for Stem Cell Engineering and Regenerative Engineering (UQ StemCARE). Bruce Conklin (Department of Medicine Gladstone Institute of Cardiovascular Disease) is greatly acknowledged for the WTC iPSCs gift. The authors acknowledge the facilities, and the scientific and technical assistance, of the Microscopy Australia Facility at the Centre for Microscopy and Microanalysis (CMM) of The University of Queensland. We thank Zoe L. Hunter for performing the recording of MEA experiments and providing the schematic diagrams. We thank the Queensland Health Forensic and Scientific Services, Queensland Department of Health, for providing SARS-CoV-2 isolate. Next Generation Sequencing was performed by ACE Sequencing (SCMB, UQ). Purchasing of NGS reagents was supported by Illumina COVID Matched Funding Scheme. We would like to unknowledge Dr Alejandro Rojas Fernandez (Universidad Austral de Chile) and Dr Alberto Amarilla (SCMB, UQ) who kindy provided W25 nanobody against SARS-CoV-2 spike protein. We are grateful to Dr Naphak Modhiran and Dr Daniel Waterson (SCMB, UQ) for providing CR3022 anti-SARS-CoV-2 spike antibody.

## AUTHOR CONTRIBUTIONS

M.S. performed, analyzed and designed experiments, interpreted the results, and wrote the manuscript, A.S. performed, analyzed and designed SARS experiments, interpreted the results, wrote the manuscript, B.A. performed additional experiments, S.D.M. performed additional experiments, J.S. performed additional experiments, J.C-W contributed to the conception of the study, A.K. contributed to the design, interpreted results, and wrote the manuscript, E.W. conceived and supervised the study, interpreted results, and wrote the manuscript. The final version of the manuscript was approved by all authors.

## DATA AVAILABILITY STATEMENT

The data that support the findings of this study are available from the corresponding author upon request. Bulk RNA-sequence data from brain organoids that support the findings of this study have been deposited in the GEO-NCBI with the primary accession codes GSE208575 and GSE208440.

## Supplemental Information

**Figure S1. Characterization of Neuroectoderm Differentiation and Spheres Formation from hPSCs, related to Figure 1**.

(**A**) Immunocytochemistry of human iPSCs stained with TRA-1-60 (Green), NANOG (Green), and SOX2 (Red). All cells were counterstained with Hoechst 33342 (Blue). Scale bar = 40 µm.

(**B**) Immunocytochemistry of human iPSCs-derived neuroectoderm labeled with NANOG (Green), SOX2 (Red), and NESTIN (Green) antibodies. All cells were counterstained with Hoechst 33342 (Blue). Scale bar = 20 µm.

(**C**) Wholemount immunostaining of hNEct spheroids at day 4 of FGF, CHIR99021, and BMP4 treatments stained with SOX2 (Red) and ZO1 (Green). The spheroid is counterstained with Hoechst 33342 (Blue). Scale bar = 100 µm.

(**D**) Wholemount immunostaining of hNEct spheroids at day 4 of FGF and different concentrations of BMP4 treatments stained with SOX2 (Green). The spheroids are counterstained with Hoechst 33342 (Blue). Scale bar = 50 µm. Right top graph represents the average intensity of SOX2 expression from center to periphery across FGF and BMP4 treated spheroids. Right bottom graph shows quantification of the percentage of cells expressing the neural stem cells marker (SOX2) relative to the total number of cells per sample. Data are presented as mean ± standard deviation. Number of independent experiments = 3. \*\*\*\**P* < 0.01 via One Way ANOVA.

(**E**) Bright-field images of CPCOs generated from hPSCs at day 28 of differentiation with and without embedding into ECM. Scale bar = 500 µm.

**Figure S2. Reproducible Organization of CPs and Cortical Cells in CPCOs Generated from Different hPSC lines, and Transcriptional Analysis of CPCOs and COs related to Figure 1**.

(**A**) Bright-field images of CPCOs generated from H9-ESCs, G22-iPSCs and WTC-iPSCs line at day 28 of differentiation. Cross indicates the organoids without emerging thin epithelium. Scale bar = 500 µm.

(**B**) Representative images of sectioned CPCOs immunostained with TTR (Green) and LMX1A (red) antibodies. All cells were counterstained with Hoechst 33342 (Blue). Scale bar = 200 µm. Dotted black lines indicates the cut bright-field CPCOs shown in (A).

(**C**) Quantification of successful CPCO generation at day 28 across different hPSC lines (H9-ESCs, G22 iPSCs and WTC iPSCs). N = 3. Data are presented as mean ± standard deviation.

(**D**) Heatmap expression of marker genes related to cortical hem, choroid plexus within the bulk RNA transcriptomes of CPCOs at days 28 and 56 and traditional “naked” COs at day 56. Values are shown as z-score.

(**E**) Heatmap expression of marker genes related to telencephalonic CP within the bulk RNA transcriptomes of CPCOs at days 28 and 56 and traditional “naked” COs at day 56. Values are shown as z-score.

(**F**) Hierarchical clustering of gene expression profiles of CPCOs at day 28 and 56, “naked” COs at day 56 and human CP tissue based on the expression of multiple CP markers. The values are read counts normalized to library sizes, composition bias and log+1-transformed.

**Figure S3. Analysis of Tight Junction and Neural Cells During the Development of CPCOs, related to Figures 2&3**.

(**A**) Heatmap expression of marker genes related to apicobasal polarity within the bulk RNA transcriptomes of CPCOs at days 28 and 56 and traditional “naked” COs at day 56. Values are shown as z-score.

(**B**) qRT-PCR of marker genes related to apicobasal polarity (*CLDN12, CLDN11,* and *PCDH18*) in CPCOs. All values were normalized to GAPDH levels of their respective samples and expressed relative to Day 7 values to obtain the fold change. Data are shown as the mean ± standard deviation; **p < 0.01, ***p < 0.001, ****p < 0.0001 via one-way ANOVA. Number of independent experiments = 3.

(**C**) Analysis of immunostaining of sections CPCOs at day 28 showing tight junction ZO1 (Green) protein expression in the epithelial cells of CPs. Section was counterstained with Hoechst 33342 (Blue). Scale bar = 72 µm.

(**D**) Analysis of immunostaining of sections CPCOs at days 56 and 150 showing oligodendrocyte progenitors marked with PDGFRA (Green), and mature oligodendrocytes marked with CNPase (Red) proteins. Section was counterstained with Hoechst 33342 (Blue). Scale bar = 40 µm.

**Figure S4. Validation of Cortical Neurons Emerging in CPCOs Generated from Different Batches, related to Figure 3**.

(**A**) Analysis of immunostained CPCO sections at day 28 obtained from different batches showing cortical neurons in layer VI marked by TBR1 (Red) and layer V marked by CTIP2 (Green) proteins. All sections were counterstained with Hoechst 33342 (Blue). Scale bar = 122 µm.

(**B**) Heatmap expression of marker genes related to oligodendroglia and astrocytes within the bulk RNA transcriptomes of CPCOs at days 28 and 56 and traditional “naked” COs at day 56. Values are shown as z-score.

(**C**) Analysis of CPCOs sections at day 28 immunostained with astrocyte marker GFAP (Red) protein. Section was counterstained with Hoechst 33342 (Blue). Scale bar = 20 µm.

(**D**) Schematic diagram summarizing the developing structure of CPCOs at different days of development showing the specified the two domains including CPs, and the cortex and the culture conditions used.

(**E**) Representative images of raster plots of a single MEA show the intensity of spiking activity before and after Bicuculline, NMDA, Glutamate treatments. Raster plots are shown in 1 min segments. E is electrode.

**Figure S5. Analysis of Euploid and Trisomy 21 CPCOs Characteristics, related to Figure 4**.

(**A**) Graph showing the growth (average diameter) of CPCOs in euploid (EU79) and trisomy 21 (DS18) at different stages of *in vitro* culture. Data are presented as mean ± standard deviation (n = 4).

(**B**) Hierarchical clustering of the transcriptomes of DS(DS18) and euploid (EU79) CPCOs based on the expression of top 500 most variable genes.

(**C**) Box blots showing distribution of differentially expression genes associated with choroid plexus and secretomes in euploid (EU79) and trisomy 21 (DS18) CPCOs at day 28 obtained from bulk RNA-seq. Data are presented as minimum to maximum, with notches are centered on the median. Values are shown as z-score.

(**D**) Heatmap expression of HSA21 genes among the top 500 most variable genes within the bulk RNA transcriptomes of euploid (EU79) and DS (DS18) CPCOs. Values are shown as z-score.

(**E**) Western blots showing the protein levels of OLIG2 in euploids (EU79 and G22) and DS (DS18 and G21) CPCOs at day 35. Actin was used for normalization. All blots derived from the same experiment and processed in parallel. Right graph shows the quantification of normalized OLIG2 level obtained from blots.

(**F**) Analysis of euploid (EU79) and DS (DS18) CPCO sections at day 35 immunostained with OLIG2 (Green) and SOX2 (Red) antibodies. All sections were counterstained with Hoechst 33342 (Blue). Scale bar = 20 µm.

(**G**) Box blot showing distribution of marker genes (listed below in the heatmap) associated with interneurons obtained from bulk RNA-seq of CPCOs at day 28 derived from euploid (EU79) and DS (DS18) lines. Data are presented as minimum to maximum, with notches centered on the median. Below heatmap expression of marker genes related to cilia. Values are shown as z-score.

**Figure S6. Analysis of Oligodendroglia, Apicobasal Polarity, and Electrophysiological Activity Characteristics of Euploid and Trisomy 21 CPCOs, related to Figures 4, 5**.

(**A**) Violin blot showing distribution of marker genes (listed below in the heatmap) associated with apicobasal polarity obtained from bulk RNA-seq of CPCOs at day 28 derived from euploid (EU79) and DS (DS18) lines. Data are presented as minimum to maximum, with individual dots represent individual replicate of genes listed below. **p < 0.01 via Student’s t-test. Below heatmap expression of marker genes related to apicobasal polarity. Values are shown as z-score.

(**B**) Box blot showing distribution of marker genes (listed below in the heatmap) associated with oligodendroglia obtained from bulk RNA-seq of CPCOs at day 28 derived from euploid (EU79) and DS (DS18) lines. Data are presented as minimum to maximum, with individual dots represent individual replicate of genes listed below. ****p < 0.0001 via Student’s t-test. Below heatmap expression of marker genes related to oligodendroglia. Values are shown as z-score.

(**C**) Western blots showing the protein levels of oligodendrocyte precursors (PDGFRA, SOX10) and cell polarity (ZO1, CDH7, E-CAD, β-CATENIN) in euploid (EU79) and DS (DS18) CPCOs at day 49. Actin was used for normalization. All blots derived from the same experiment and processed in parallel.

(**D**) Analysis of euploid (EU79) and DS (DS18) CPCO sections at day 56 immunostained SOX10 (Red) and the CP marker TTR (Green) antibodies. All sections were counterstained with Hoechst 33342 (Blue). Scale bar = 30 µm.

(**E**) Bar graph showing the percentage of SOX10 expressing cells in euploid (EU79) and DS (DS18) CPCOs at day 56. Data are presented as mean ± standard deviation. *p < 0.05 indicates statistical significance via Student’s t-test. Number of independent experiments = 3.

(**F**) Quantification of cilia length in CPCOs derived from euploid (EU79) and DS (DS18) lines at day 56 of differentiation. Data are presented as mean ± standard deviation; ns is not significant via Student’s t-test. Number of independent experiments = 3. Individual dots represent a cilium length.

(**G**) Box blot showing distribution of marker genes (listed below in the heatmap) associated with neural cells obtained from bulk RNA-seq of CPCOs at day 28 derived from euploid (EU79) and DS (DS18) lines. Data are presented as minimum to maximum, with individual dots represent individual replicate of genes listed below. ****p < 0.0001 via Student’s t-test. Below heatmap expression of marker genes related to neural cells. Values are shown as z-score.

(**H**) Representative images of raster plots of a single MEA show the intensity of spiking activity in CPCOs at day 28 derived from euploid (EU79) and DS (DS18) lines. before and after Bicuculline, NMDA, Glutamate treatments. Raster plots are shown in 1 min segments.

(**I**) Bar graph showing the mean bursting rate in neural activity in CPCOs at days 28 derived from euploid (EU79) and DS (DS18) lines. Bar graphs represent mean ± SD in firing rate, *p < 0.05 via Student’s t-test.

(**J**) Bar graphs showing the changes in the patterns of mean firing rate before and after drug treatments. The following drugs were used: 50 µM Glutamate, 10 µM NMDAA, and 50 µM Bicuculline. *p < 0.05, **p < 0.01, ***p < 0.001, ****p < 0.0001 via Two-way ANOVA.

(**K**) Representative image of trisomy 21-derived CPCOs at day 35 immunostained with ACE2 (Red), and TTR (Green). Section was counterstained with Hoechst 33342 (Blue). Scale bar = 100 µm.

(**L**) Bar graphs showing the amount of SARS-CoV-2 related genes *Envelop* and *Spike* in euploid (EU79) and trisomy 21 (DS18) CPCOs at day 28. Data are presented as mean ± standard deviation. **p < 0.01, ****p < 0.0001 via 2-Way ANOVA. Number of independent experiments = 3.

**Figure S7. Expression of SARS-CoV-2 related genes in Organoids, related to Figure 6**.

(**A**) Hierarchical clustering heat map of the top 500 most variable genes expressed in DS18 and EU79 CPCOs that were infected with SARS-CoV-2 or uninfected.

(**B**) Expression of host genes required for SARS-CoV-2 infection in DS18 and EU79 CPCOs. The values represent normalized read counts expressed as counts per million (cpm). The graphs show the mean values from 3 biological replicates with error bars indicating SDs. The statistical analysis was performed by Student’s t-test. ***P<0.001.

(**C**) Gene ontology (GO) enrichment analysis of differentially expressed genes identified in Figure 6F. Z-scores indicate the cumulative increase or decrease in expression of the genes associated with each term. Size of the bubbles is proportional to the number of DEGs associated with respective GO term. MF – molecular function, CC – cellular component.

(**D-E**) Representative confocal images of day 31 CPCOs derived from euploid (EU79) and DS (DS18) immunostained with NEUN (Green) and SARS-CoV-2 spike (Red) after SARS-CoV-2 (10^6^ FFUs). All sections were counterstained with Hoechst 33342 (Blue). Scale bar = 10 µm.

**Figure S8. Analysis of Apoptotic Process in SARS-CoV-2 infected CPCOs, related to Figure 5&7**.

(**A**) Violin blot showing distribution of marker genes (listed below in the heatmap) associated with apoptosis obtained from bulk RNA-seq of day 31 CPCOs before and after SARS-CoV-2 infection derived from euploid (EU79) and DS (DS18) lines. Data are presented as minimum to maximum, with individual dots represent individual replicate of genes listed below. **p < 0.01; ****p < 0.0001 via Student’s t-test. Below heatmap expression of marker genes related to apoptosis. Values are shown as z-score.

(**B**) Analysis of sections of euploid (EU79) and DS (DS18) day 31 CPCO infected with Mock and SARS-CoV-2 over 72 hpi immunostained with Cleaved caspase-3 (Red) and SARS-CoV-2 spike (Green) antibodies. All sections were counterstained with Hoechst 33342 (Blue). Scale bar = 35 µm. Below violin graph showing the percentage of cleaved caspase 3 positive cells in CPCOs derived from euploid (EU79) and DS (DS18) infected with the Mock and SARS-CoV-2 virus. **p < 0.01; ***p < 0.001; ****p < 0.0001 via 2-Way ANOVA.

(**C**) Quantification of viral titers from CPCOs culture supernatants of euploid (EU79), DS (DS18), and EU79 treated with 100uM avoralstat or camostat or nafamostat after SARS-CoV-2 (10^6^ FFUs) infection of day 28 CPCOs at 0, 24, and 48 hpi. Data are presented as mean ± SEM. N = 3 biological replicates consisting of 3 organoids each; **p < 0.01; ****p < 0.0001 via One-Way ANOVA.

