## Supplementary figures and images for "Choroid plexus defects in Down syndrome brain organoids enhance neurotropism of SARS-CoV-2"

### Supplementary Figures

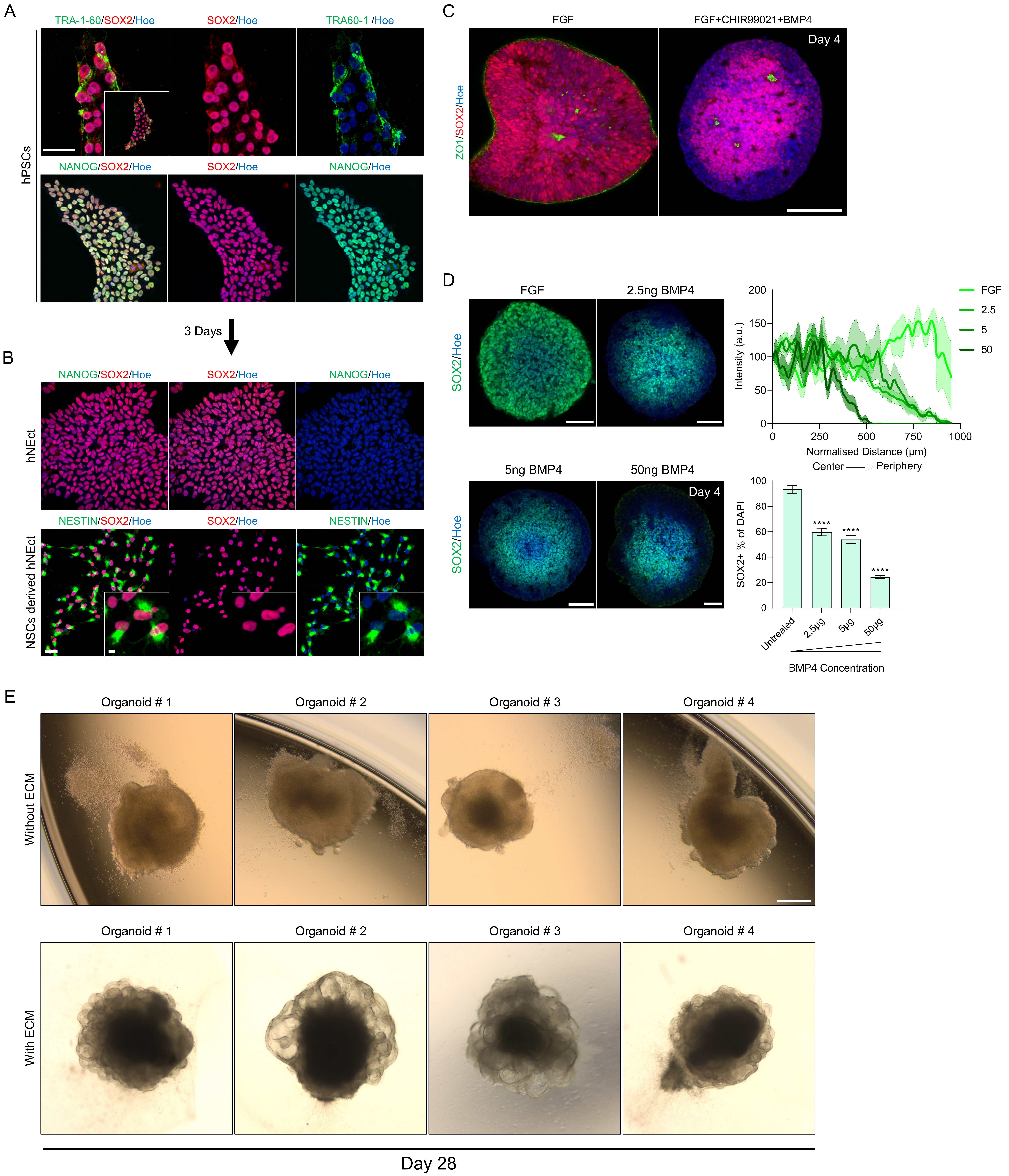

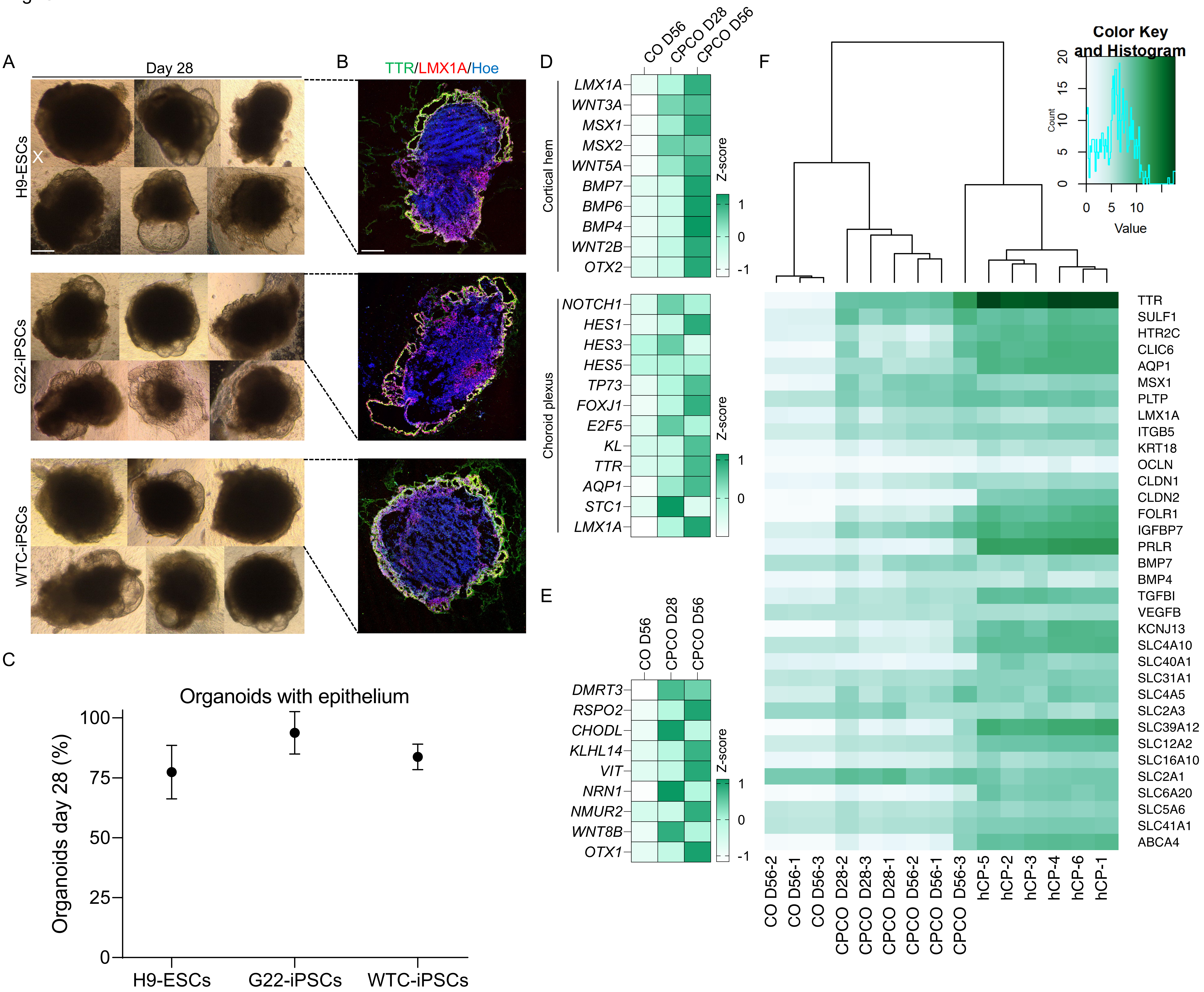

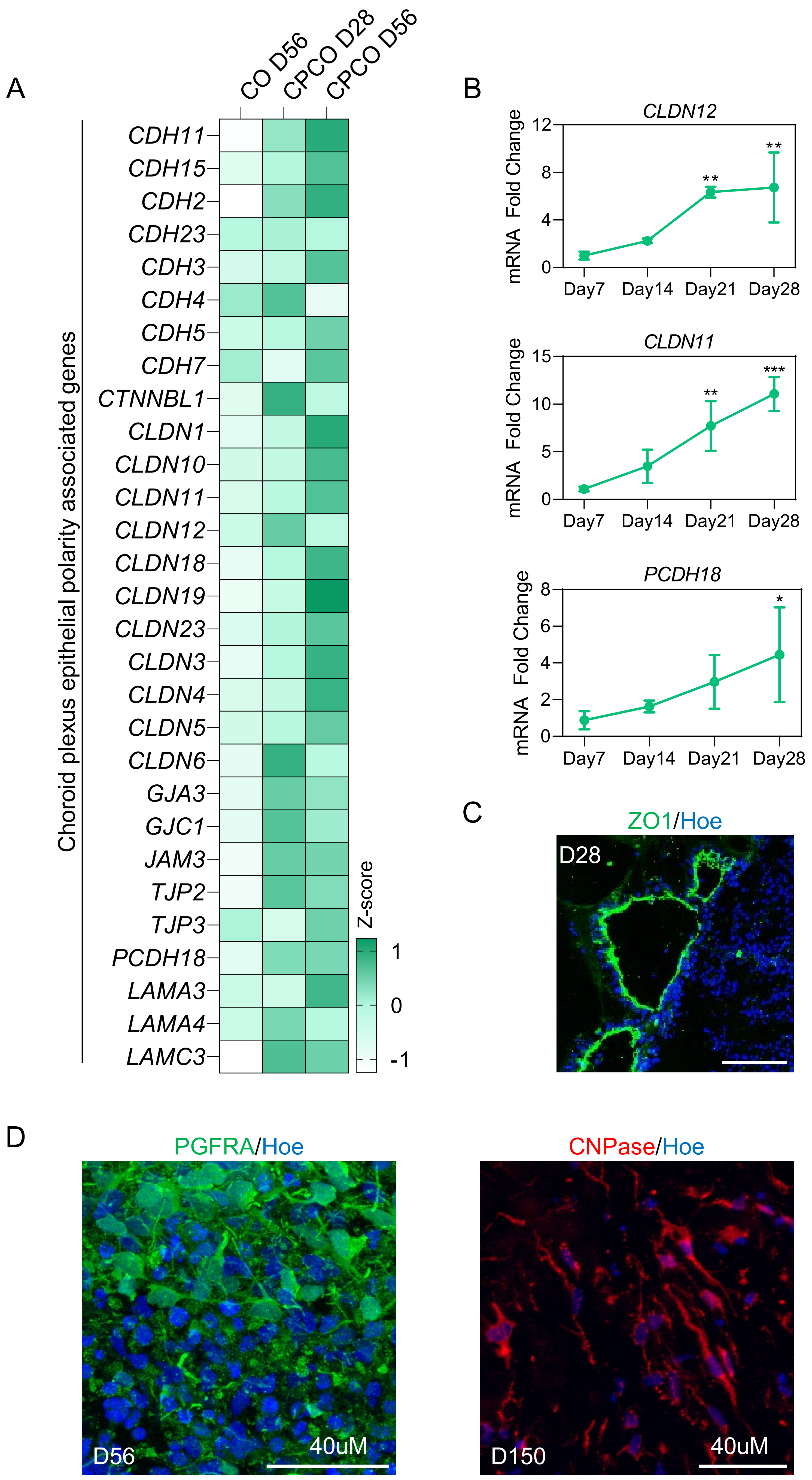

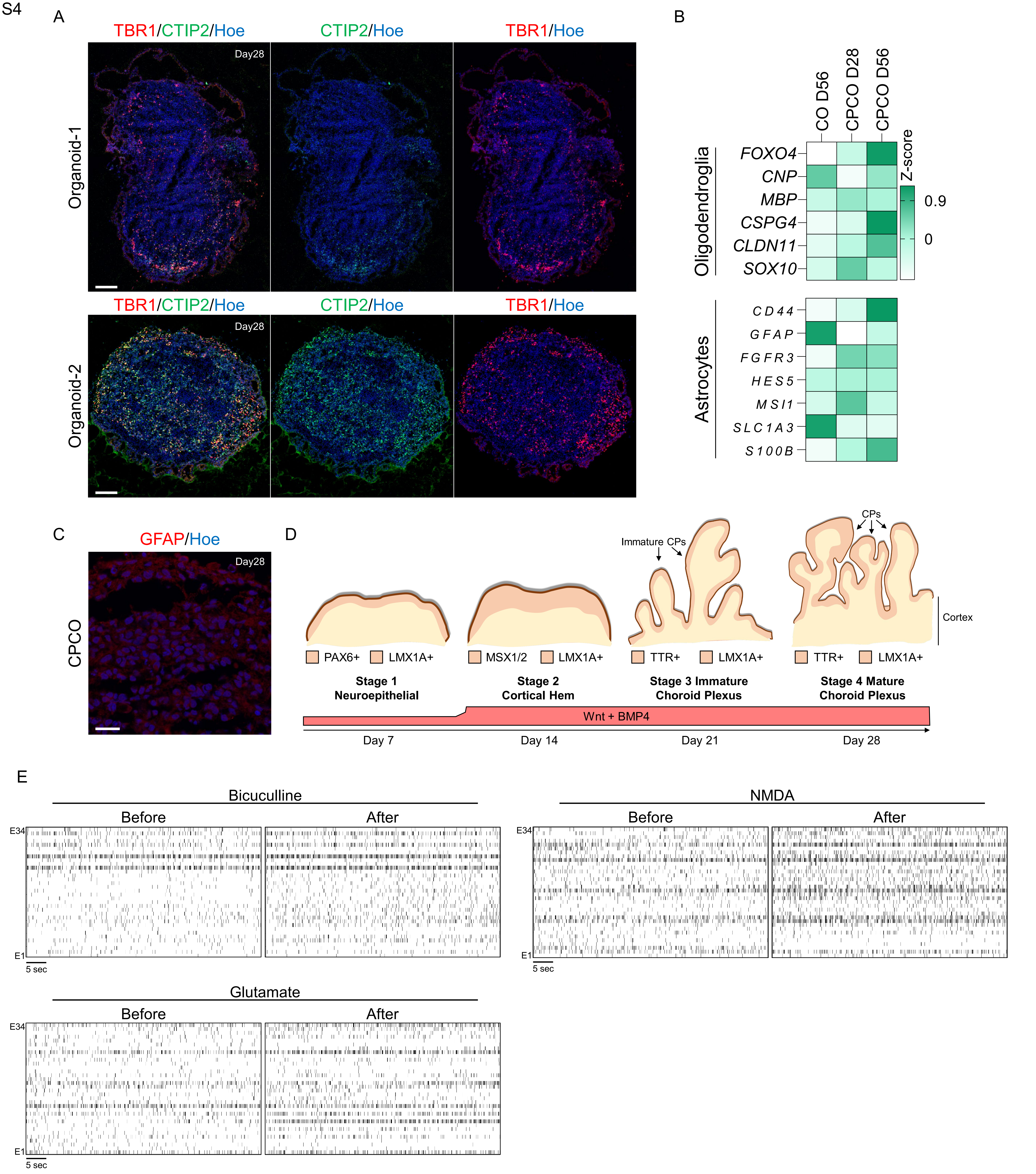

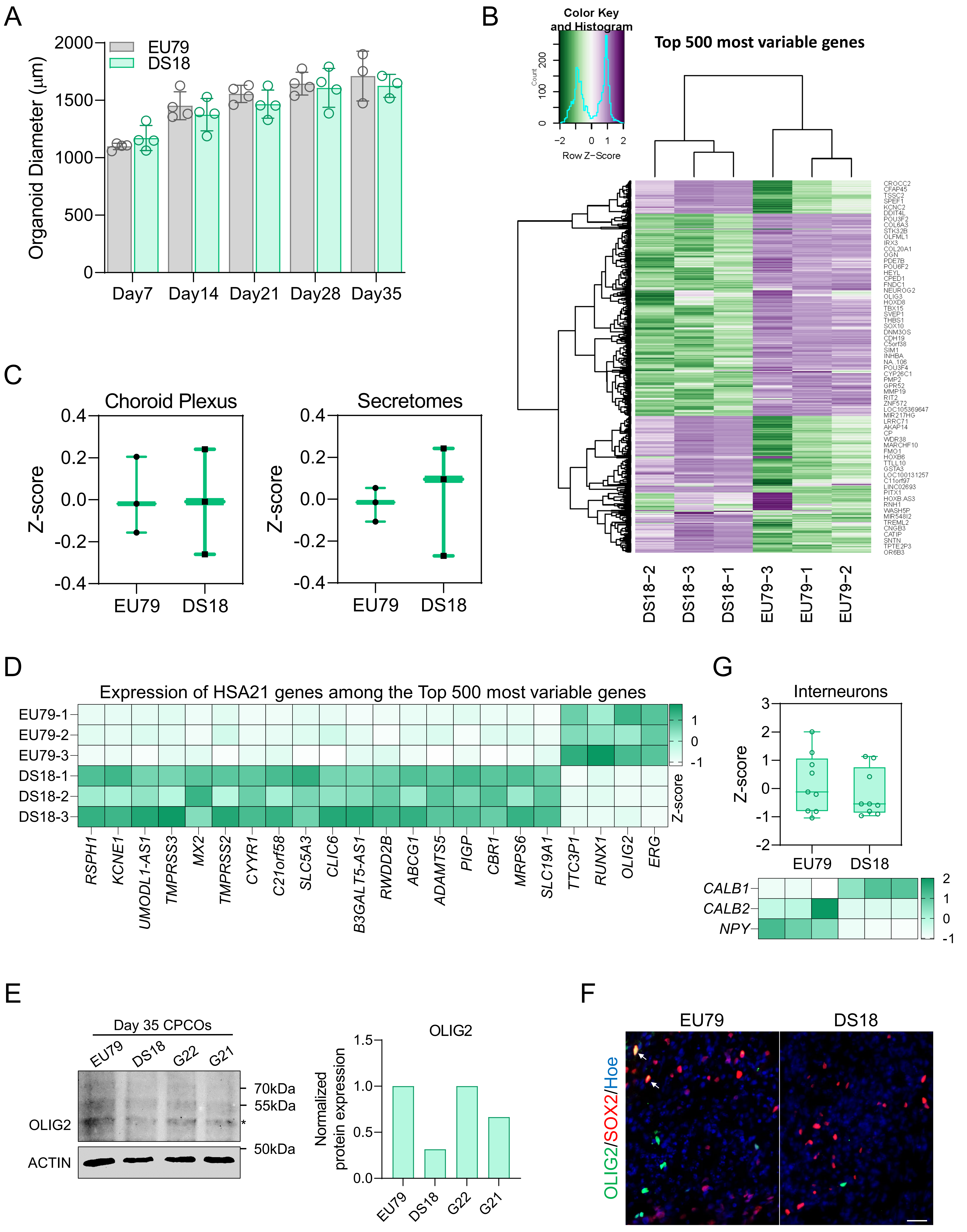

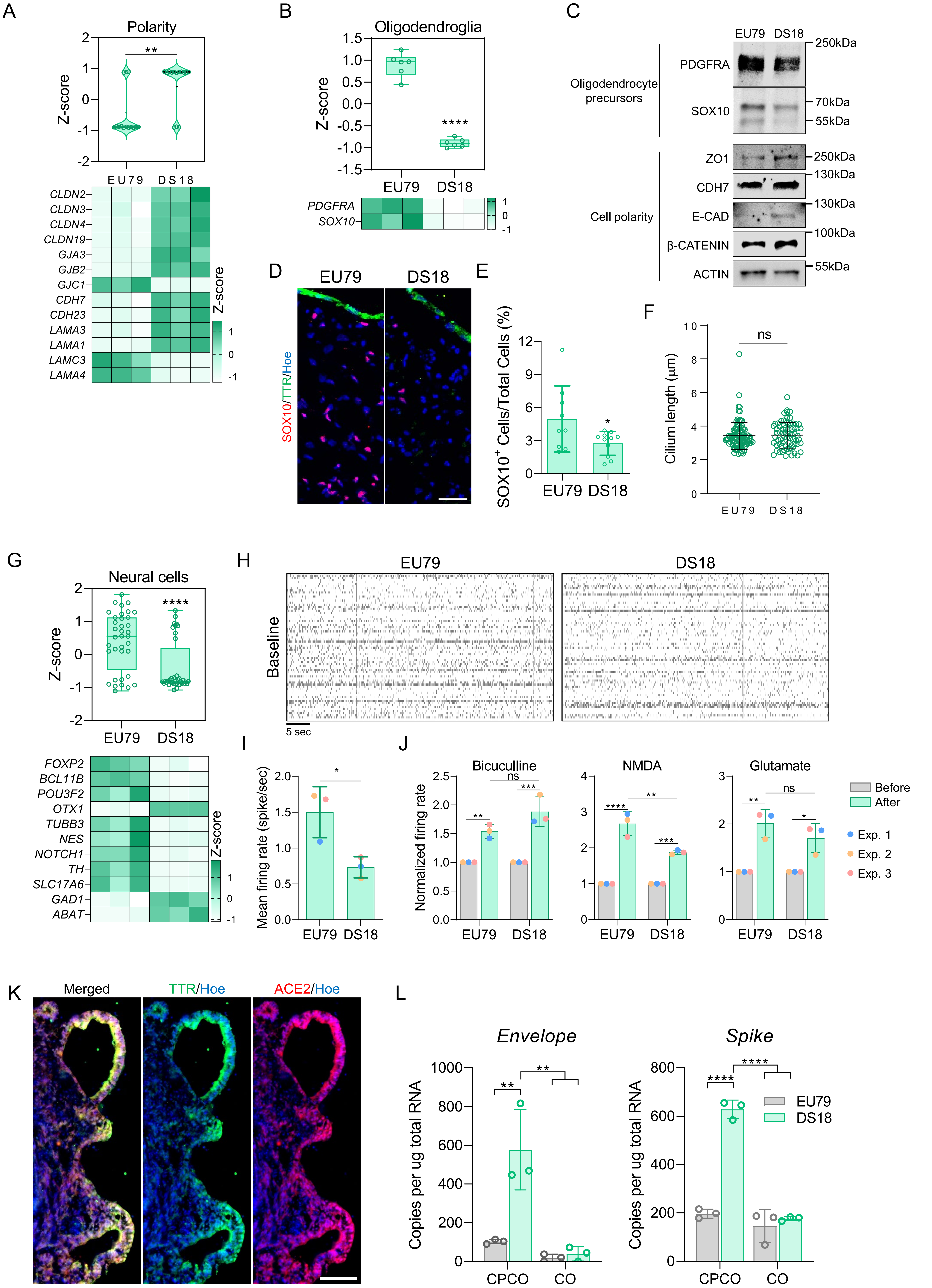

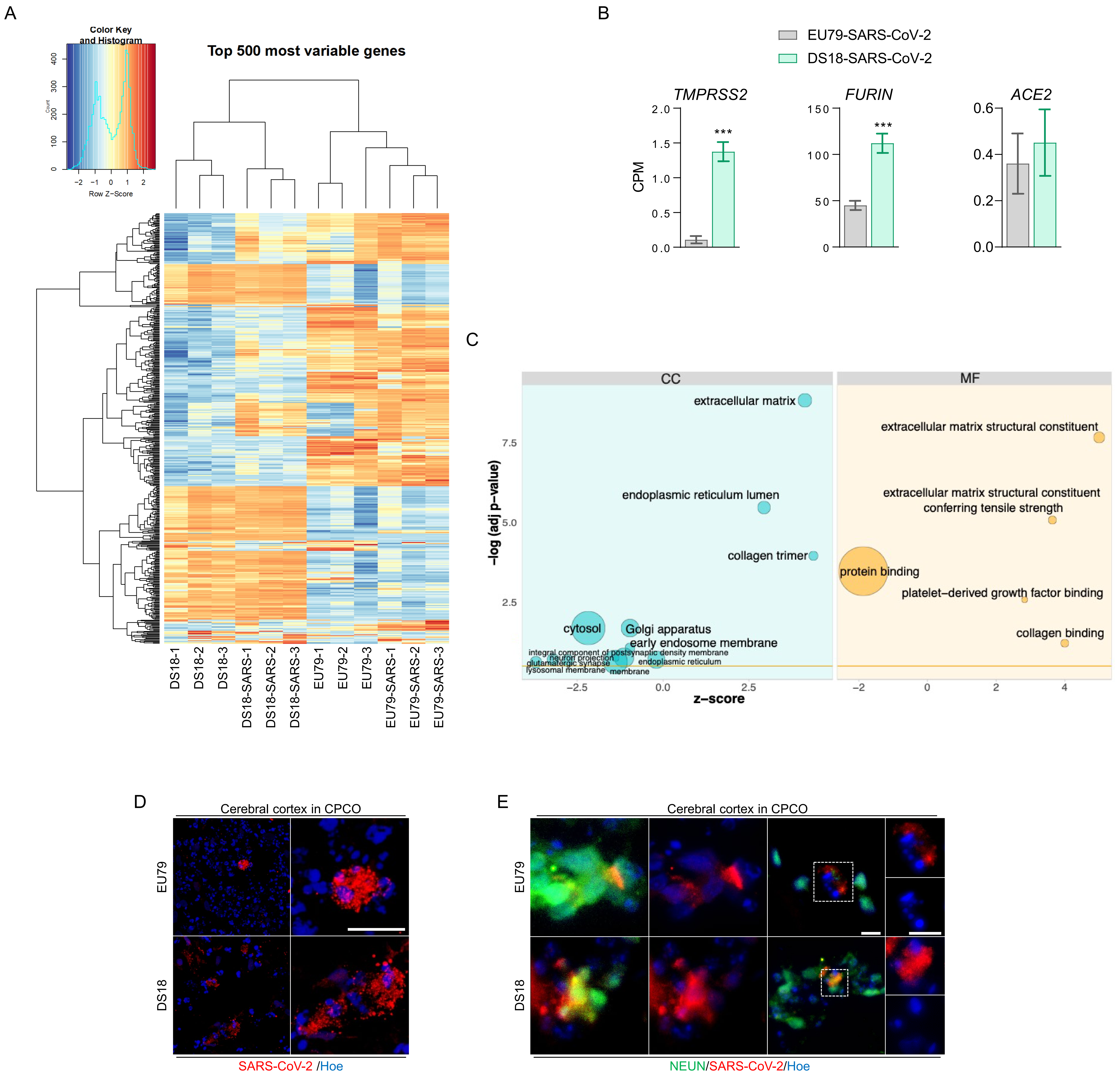

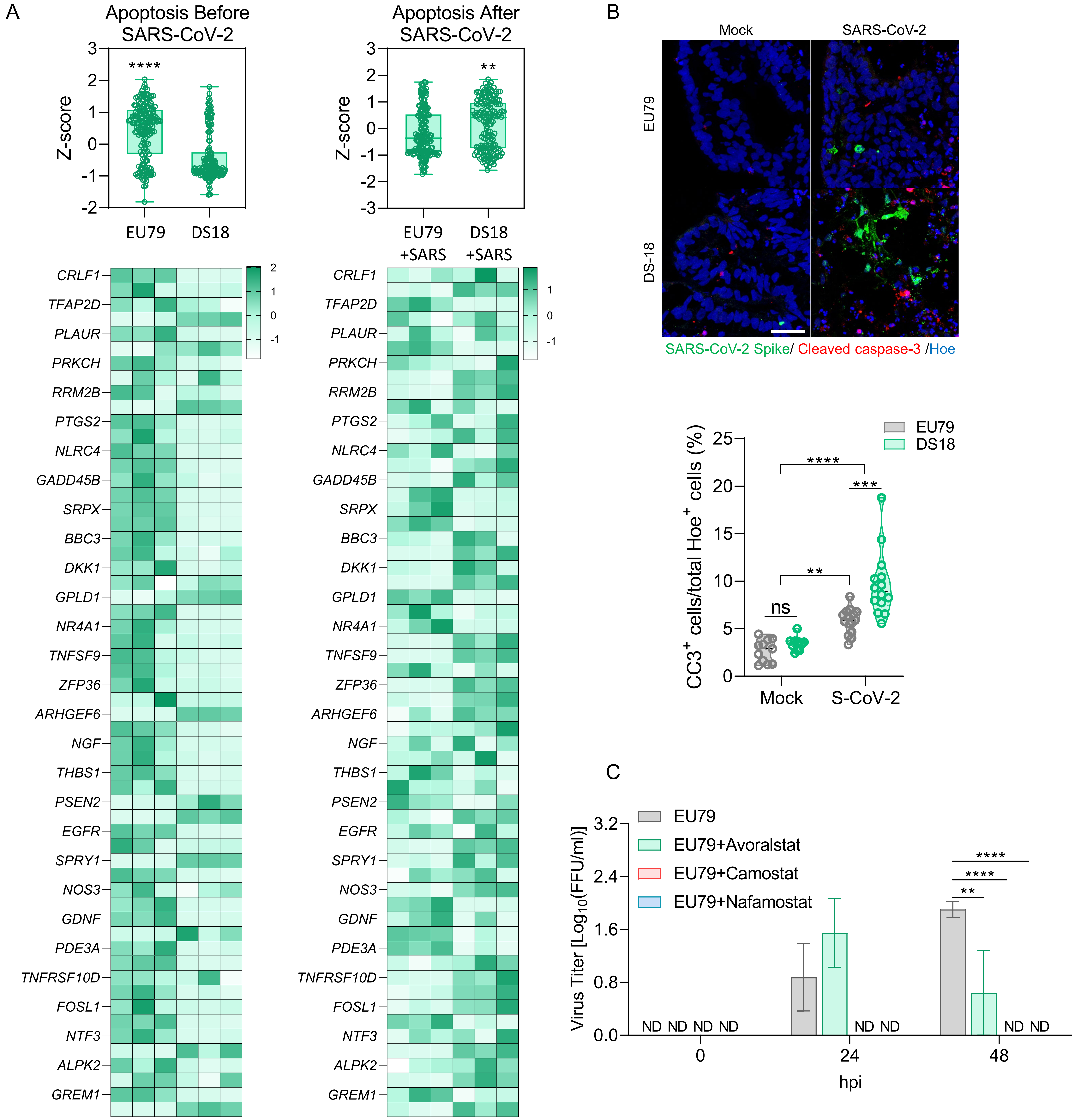
